# A learned fungal ITS embedding does not outperform correctly configured alignment in open-world evaluation

**DOI:** 10.64898/2026.09.17.752437

**Authors:** Aaron O’Brien, Auguste Gardette

## Abstract

1. Learned sequence embeddings are increasingly proposed for fungal ITS classification and novelty detection, but are usually evaluated against weaker versions of themselves or default-configured alignment, by AUROC alone, and with queries treated as independent. We asked whether a purpose-trained encoder outperforms percent identity when both score identical queries under a firewalled open-world design, and which evaluation choices decide the answer.
2. From the UNITE release of 19 February 2025 we built ITS-core, ITS1 and ITS2 views and a genus-separated split with sealed calibration and test partitions. A convolutional encoder trained with genus-proxy, cross-view, hierarchical and episodic objectives was frozen and hash-sealed before test data were opened. We compared it with exhaustive, coverage-filtered VSEARCH identity on the same queries and references using genus-clustered paired intervals, and logged every post-opening correction before computing corrected metrics.
3. Three evaluation choices changed the comparison. Default VSEARCH heuristics returned a lower-identity hit than exhaustive search for 48.3% of benchmark queries; without a coverage filter, identity placed only 64.9% of novel ITS-core queries in the correct class, against 98.7% with it, because the conserved 5.8S let partial alignments win; and the encoder’s embeddings depended on inference batching. Once corrected, identity exceeded the encoder in known-genus accuracy and novel-family placement at every view (family 80.0% to 84.6% against 53.8% to 64.8%) and detected more novel genera at a lower false-novelty rate. The encoder’s novelty AUROC was within 0.023 of identity’s, and no genus-clustered interval excluded zero. Identity’s own development-to-test gap exceeded the encoder’s, so that gap reflects partition composition rather than selection. Of the genera held out by a historical benchmark, 85% had been present in training; on a leakage-safe version, identity’s AUROC advantage was 0.065, with a genus-clustered interval excluding zero.
4. Correctly configured alignment matched or exceeded the learned embedding throughout. The decisive results came from the evaluation, not the representation: baseline configuration, batch invariance, genus-level uncertainty, a parameter-free control for selection and a leakage audit each changed a conclusion, and each is inexpensive to apply to any learned barcode method.

## 1. Introduction

Fungal metabarcoding routinely returns sequences whose closest named relative is distant and whose genus is absent from every reference collection [9]. A method deployed on such data has two jobs: to decide whether a query is novel enough to withhold a genus assignment, and to place it as precisely as the reference allows. Learned sequence embeddings are an attractive answer to both, because they promise a single similarity space in which unseen genera sit near their relatives and in which different barcode views become comparable [2, 3, 13].

How such methods are evaluated matters as much as how they are built. Four habits are common. Learned models are compared chiefly with weaker variants of themselves, so an ablation ladder can show steady improvement without saying whether any rung beats a method practitioners already use. When an alignment baseline is included it is usually run with default settings. Novelty is summarised by AUROC, a whole-curve statistic, rather than by the operating points at which a user would actually flag sequences. And uncertainty is computed over queries as if they were independent, although queries from one genus share most of their information. A recent systematic comparison of classifiers for unknown barcodes, including fungal ITS, found composition-based classifiers strongest and showed that apparent under-classification by alignment often reflected threshold stringency rather than genuine novelty [25]; baseline configuration, in other words, can decide the result.

We built a purpose-trained ITS encoder under a deliberately strict design: a genus-separated split, a controlled ladder that adds one training objective at a time, a hash-sealed freeze before calibration and test data were opened, and split-conformal calibration of the novelty decision [10]. The sealed evaluation supported a positive claim about higher-rank placement and cross-view retrieval. It did not include a non-learned baseline on the same split. Adding one, and examining the evaluation that produced the claim, is the subject of this paper.

The contribution is therefore an evaluation protocol with the encoder as its worked example. We report five checks, each of which changed a conclusion here: auditing the alignment baseline’s search configuration; testing whether embeddings are invariant to inference batching; computing paired uncertainty at the genus rather than the query level; separating selection optimism from partition composition with a parameter-free control; and auditing a benchmark for training-set leakage before scoring it. Under the corrected evaluation, a correctly configured alignment baseline matched or exceeded the learned embedding on every comparison we could make.

## 2. Methods

### 2.1 UNITE collection and ITS views

We used the UNITE dynamic general release dated 19 February 2025 [9, 17]. The source FASTA contained 102,137 records. ITS regions were extracted with ITSx [4], run on the standard trimmed file (`sh_general_release_dynamic_19.02.2025.Fasta`) rather than its `_dev` variant, which holds the same records; the input lengths ITSx recorded matched the trimmed file for 500 of 500 sampled records and the `_dev` file for 323. The historical ITS2 collection used by the companion novelty workflow contains 99,808 sequences; a fresh extraction recovered 99,862. Every historical record was contained in the fresh extraction, so the difference is 54 additional calls rather than any loss of historical sequences.

Three views were retained where annotated: ITS1, ITS2, and *ITS-core*, the contiguous ITS1— 5.8S—ITS2 span. Exact sequence hashes identified 301 view-level groups in which an identical ITS1 or ITS2 sequence carried more than one genus label. Rather than discard the whole source record, we masked only the conflicting view: 393 ITS1 views and 338 ITS2 views. No cross-genus exact conflict was detected for ITS-core. Requiring a named family and genus left 67,406 records for open-world splitting (Table 1).

**Table 1:** Source collection and open-world partitioning. *Records* counts source records after taxonomic filtering; the three right-hand columns give how many of those records carry each barcode view and are therefore the denominators for the per-view results in Tables 5–7.

| item | records | records carrying the view |  |  |
| --- | --- | --- | --- | --- |
|  |  | ITS-core | ITS1 | ITS2 |
| *Source collection* |  |  |  |  |
| UNITE dynamic release (19 Feb 2025) | 102,137 | — | — | — |
| historical ITS2 pool | 99,808 | — | — | — |
| fresh ITS2 extraction | 99,862 | — | — | — |
| named family and genus | 67,406 | — | — | — |
| *Open-world partitions* |  |  |  |  |
| TRAIN_REF | 46,453 | 46,151 | 46,342 | 46,076 |
| DEV_KNOWN | 1,528 | 1,515 | 1,527 | 1,515 |
| DEV_NOVEL | 8,698 | 8,659 | 8,682 | 8,642 |
| CAL_KNOWN | 1,505 | 1,494 | 1,501 | 1,493 |
| TEST_KNOWN | 1,484 | 1,467 | 1,477 | 1,463 |
| TEST_NOVEL | 7,738 | 7,718 | 7,702 | 7,671 |
| total partitioned | 67,406 |  |  |  |

Exact cross-genus sequence conflicts were masked at the level of the individual view rather than by deleting the source record: 393 ITS1 and 338 ITS2 views across 301 conflict groups. No ITS-core conflict was detected.
The six partitions sum exactly to the named-family-and-genus total.

### 2.2 Open-world split and evaluation firewall

The split was built so that the genera defining each novel population were absent from the corresponding reference partition while their higher-rank lineage remained represented. Placeholder genus and species labels were not allowed to define held-out taxa. TRAIN_REF contained 46,453 records and was the only partition to receive gradient updates; the remaining five partitions are given in Table 1 and sum exactly to the filtered total. DEV_KNOWN and DEV_NOVEL drove model selection. CAL_KNOWN, TEST_KNOWN and TEST_NOVEL were excluded from all tuning (Figure 1a).

**Figure 1:**
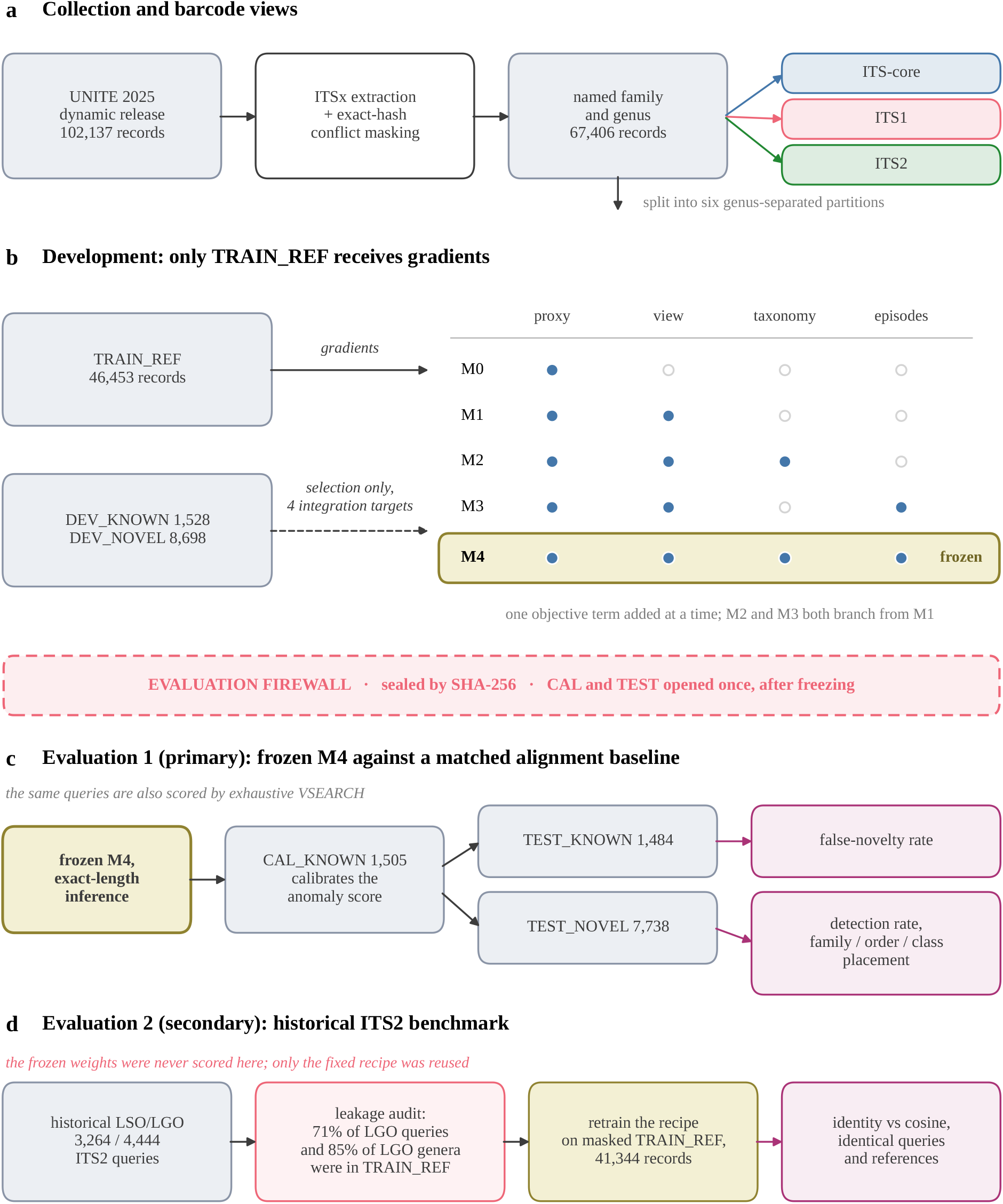
Design of the study. Fill colour marks the kind of object: grey for a data partition, white for a process, yellow for a model, purple for a reported result. (a) Collection, conflict masking and the three barcode views. (b) Development: only TRAIN_REF receives gradients, and the matrix gives the objective terms of each rung (Appendix B). (c) The primary evaluation, run once after freezing, in which the same queries are scored by the frozen encoder with exact-length inference and by exhaustive VSEARCH identity. (d) The historical benchmark: most of its held-out taxa had entered the original training split, so the frozen weights were not scored there and the same recipe was retrained on a masked TRAIN_REF. The dashed band separates what was iterated on from what was opened once.

After M0–M3 had been examined, but before M4 was trained, we fixed four integration targets. Each target was set at or slightly below the best value already achieved by a specialized rung, so that M4 would be required to preserve both novelty discrimination and higher-rank placement rather than optimize either endpoint alone. The targets were ITS2 AUROC ≥ 0.73,ITS2 novel-family placement ≥ 40%, ITS-core AUROC ≥ 0.74, and ITS-core novel-family placement ≥ 50%. Had any target been missed, M4 would not have replaced the best specialized model as the final candidate. Under the padded inference then in use, M4 achieved 0.751, 41.3%, 0.778 and 57.5%, respectively. Rescored with exact-length inference after the rescore had been pre-registered (Section 2.6), it achieved 0.754, 40.3%, 0.777 and 57.6%, so it still met all four targets, the ITS2 placement target by 0.3 points. We record the standing of this pre-specification precisely, since it bears on how the frozen evaluation should be read. The deposited run logs date every M0–M3 fit before the M4 fit, so the order of the experiments is checkable from the artefacts; the four threshold values themselves were fixed in working notes in the interval between those fits and carry no independent timestamp. We therefore report them as an asserted pre-specification rather than a registered one. The selected checkpoint and split table were then sealed by SHA-256 before CAL and TEST were evaluated. The recorded hashes are in Appendix A, and the locked evaluator refused to run if either changed.

### 2.3 Encoder, retrieval score and inference

Sequences were one-hot encoded as four nucleotide channels; ambiguous bases were represented by all-zero columns, and no fixed sequence-length truncation was applied. The encoder begins with a 1D convolution (4 → 64 channels, kernel 9), followed by 12 residual convolutional blocks. Dilations 1, 2, 4, 8, 16, 32 are repeated twice; each block uses group normalisation (8 groups), GELU activations, two kernel-5 convolutions at the same dilation, and 0.1 dropout. A final group normalisation is followed by masked global mean and maximum pooling. Their concatenation is projected by a two-layer MLP (128 → 256 → 256, GELU, 0.1 dropout) and L2-normalised, producing *z* = *f*_*θ*_(*x*)/∥*f*_*θ*_(*x*)∥ . The encoder contains 597,440 trainable parameters; the 4,231-class genus-proxy head used in training adds 1,083,136. The training objectives and the development ladder are described in Appendix B.

During training and in the sealed evaluation, batches were padded to the longest sequence in the batch. Padded positions were excluded from the final pooling, but not from the group-normalisation statistics or from the receptive fields of the dilated convolutions, so an embedding computed in a padded batch depends on the other sequences in that batch (Section 3.1). Except where results are labelled as sealed, embeddings were computed with exact-length batching, in which every inference batch holds sequences of a single length, in float32 with autocast and TF32 disabled. Each embedding is then a function of its own sequence alone. The model weights are unchanged: the correction applies to inference only, to the hash-verified checkpoint.

The primary similarity for query *q* is its closest same-view reference embedding,

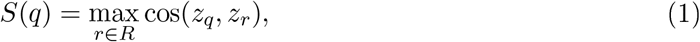

where *R* is the corresponding TRAIN_REF view. Novelty uses *A*(*q*) = −*S*(*q*), so larger values are more anomalous. Nearest-neighbour taxonomy is read from the reference attaining the maximum cosine.

### 2.4 Alignment baseline

Percent identity was computed with VSEARCH v2.31.0 [14]. VSEARCH ranks database sequences by the number of *k*-mers they share with the query and aligns candidates in that order. Under the default `-maxaccepts 1` it returns the first candidate that satisfies -id; at -id 0. 5 that is usually the top *k*-mer candidate rather than the highest-identity target, so `--top_hits_only` has no eflect and the reported value is not a best-hit identity. The default `-maxrej ects 32` also abandons a divergent query after 32 rejected candidates. The sealed historical analysis used these defaults. Every other identity score reported here used exhaustive search with a coverage requirement:

~~~
vsearch —usearch_global queries. fasta —db refs. fasta —id 0. 5
—maxaccepts 0 —maxrej ects 0 —top_hits_only —query_cov 0. 8
—threads 32 —userout hits. tsv —userfields query+target+id
~~~

where 0 disables both termination heuristics. Among tied best hits the first reported was kept. Depending on view, 1.5% to 8.1% of novel queries had more than one tied best hit. Bracketing the tie rule, by counting a placement correct only if every tied hit agreed with the query’s lineage (the conservative, lineage-aware reading) or if any did, changed novel-genus placement by at most 0.2 percentage points at any view and rank, so the choice of rule affects no result. Queries with no hit at identity ≥ 0.5 and query coverage ≥ 0.8 were retained, assigned a score of 49.999% for ranking, and counted as placement failures. VSEARCH discarded five ITS1 and four ITS2 reference sequences shorter than its 32-nt minimum. The coverage threshold was chosen after the unfiltered TEST placement results had been seen, so we report the full threshold sweep rather than a single value (Section 3.2).

The alignment baseline is paired with the encoder throughout: identical query sets, identical reference collections, and the same split-conformal procedure, with each score calibrated on its own CAL_KNOWN values. For cross-view retrieval, ITS1 or ITS2 queries were searched against the ITS-core references, which contain both spacers, so an alignment can find the homologous region directly.

### 2.5 Paired uncertainty

AUROC differences were tested with the DeLong method [22] at the query level. Because queries from one genus are strongly dependent, we also used a paired genus-cluster bootstrap: known and novel genera were resampled separately with replacement, every query in a sampled genus was carried at the same weight, both scores received identical weights, and the weighted Mann– Whitney AUROC was recomputed from scratch in each of 10,000 replicates. Conformal decision rates were resampled in the same way, holding the calibration scores fixed. Point estimates are query-weighted.

### 2.6 Post-opening corrections and deviation log

Three corrections were made after the sealed evaluation had been opened: the exact-length inference described above, the exhaustive VSEARCH configuration, and the coverage filter. Each was recorded in a version-controlled deviation log before any corrected metric was computed. The first entry was committed at 12:15 on 22 September 2026 (log SHA-256 `f49548118ec4262a4dff430a68591b226efef0ee2daf4d72f153b9fa50ef6faf`), and the first corrected TEST metrics were written at 12:33. A second entry, pre-registering how a development rescore would be reported if M4 no longer met its integration targets, was committed at 17:08 (SHA-256 `ff80854b923f3963adfc07f3a791b0199b37cc285cf2d333aafb7e32ea844f73`), one minute before the corrected development metrics were written. These timestamps come from the authors’ own machine and are not independently attested. No model was retrained, no threshold was tuned on TEST, and the sealed results are reported beside the corrected ones throughout.

### 2.7 Frozen conformal evaluation

After M4 was frozen, CAL_KNOWN alone calibrated the anomaly score. For a query with anomaly *A*(*q*) and calibration anomalies 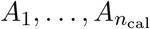, the split-conformal *p*-value is

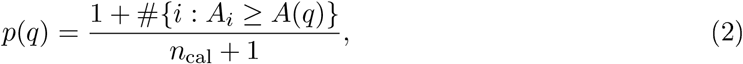

and queries with *p* ≤ *α* are called novel [19, 1]. Under exchangeability of CAL_KNOWN and a future known query, this conformal *p*-value is super-uniform, so the marginal probability of a false novelty call is at most *α*. The empirical false-novelty fraction on a finite TEST_KNOWN sample may lie above or below *α*. We report *α* ∈ {0.01, 0.05, 0.10, 0.20}. TEST_KNOWN supplies the observed false-novelty rate and TEST_NOVEL the detection rate, both per query and averaged within novel genus. No DEV sequence was embedded or scored during the locked evaluation.

### 2.8 Historical benchmark and leakage audit

The secondary benchmark uses the deposited leave-one-species-out (LSO) and leave-one-genus-out (LGO) workflow of the companion novelty study [10]. On the historical 99,808-sequence ITS2 FASTA, the currently committed splitter generates 3,264 LSO queries whose genus remains represented and 4,444 LGO queries whose family remains represented. The LGO count reproduces the earlier analysis exactly. The earlier manuscript reported 3,170 LSO queries; the repository contains only one committed splitter implementation, so the origin of that 94-query difference cannot be reconstructed. We therefore treat the present 3,264*/*4,444 split as a new paired reproduction rather than a numerical reproduction of the earlier LSO run.

Before any neural scoring we audited whether the historical held-out taxa had appeared in M4 training. They had, extensively: 3,166 of 4,444 exact LGO queries, and 458 of the 538 held-out LGO genera, were present in the original TRAIN_REF (Table S2; Figure S1). We therefore did not score the frozen M4 checkpoint on this benchmark. Instead, holding architecture and hyperparameters fixed, we removed every historical LGO query genus and every historical LSO query species from TRAIN_REF and retrained the same recipe without opening DEV, CAL or TEST. The mask removed 5,109 records (46,453 → 41,344); after view-availability filtering the training runner used 41,065 paired records covering 3,748 genera, compared with 46,150 pairs and 4,231 genera in the primary M4 run. The leakage-safe model is therefore fitted to 11% fewer records and 483 fewer genera than primary M4. Because DEV was not reopened for this run, how much that mask cost in model quality is unmeasured. That is a deliberate price for keeping the firewall closed rather than an oversight, but it qualifies the magnitude of the comparison in Table S6 and is carried forward accordingly.

Identity on this benchmark used the same exhaustive, coverage-filtered search as the primary evaluation (Section 2.4), and cosine used exact-length inference on the leakage-safe checkpoint. The sealed analysis, run with the default-settings search and padded inference, is deposited as `data/historical_benchmark_sealed.csv`; correcting both arms moved the AUROC difference from +0.069 to +0.065. VSEARCH discarded five reference sequences shorter than its 32-nt minimum in each benchmark search. Thirty LSO and 619 LGO queries had no hit at identity ≥ 0.5 and query coverage ≥ 0.8 (21 and 542 under the default-settings search) ; these queries were retained and assigned a score of 49.999% for ranking, representing censoring below the search threshold rather than an estimated identity. M4 scored every query. The same half of the LSO queries (*n* = 1,632), drawn by `np. Random. default_rng(0).permutation`, calibrated both scores; the other 1,632 served as known test cases, and all 4,444 LGO queries served as the novel population. The halving determines only which knowns calibrate and which are scored, but the operating points depend on it: across ten halvings (seeds 0 to 9) the identity advantage ranged from +0.063 to +0.075 AUROC, and every genus-clustered interval excluded zero. Seed 0, reported throughout, lies near the low end of that range.

The known population in this benchmark is therefore the LSO query set, whose species has been removed from the reference while its genus is retained. It is species-novel rather than fully known, unlike TEST_KNOWN in the primary evaluation, which shifts the conformal null distribution and makes the absolute operating points of the two evaluations non-comparable. The identity-versus-cosine contrast within this benchmark is unaffected, because both scores are calibrated and tested on exactly the same populations.

Because the deposited harness selects LSO and LGO queries under independent protocols, the two populations are not disjoint. Of the 7,708 query rows, 336 are sequences that appear in both arms; 182 of these calibrate the conformal threshold and also appear among the novel queries, and 281 genera are represented in both classes. Both scores are affected identically, so the paired contrast is unaffected, but the conformal operating points on this benchmark should be read as approximate.

## 3 Results

### 3.1 The sealed evaluation reproduced exactly but was not batch-invariant

Rerunning the locked evaluator from the hash-verified checkpoint reproduced every sealed value exactly. A second scoring script applied to the same checkpoint and queries did not: maximum cosine similarities differed by a mean of 0.005 (maximum 0.115), and by more than 10^−3^ for 6,683 of 10,679 ITS-core queries, although the most discrepant input sequences were identical (Table 2). The cause was padding. Scoring TEST_KNOWN ITS-core queries one at a time, with no padding, moved maximum cosine by a mean of 0.022 relative to padded batches of 64, and padded values were lower by 0.011 on average; batches of 64 and 256, both heavily padded, differed by only 0.004. Between unpadded and padded scoring, 35.2% of known queries changed their nearest reference and 16.6% changed its genus. Exact reproduction had shown that the evaluation was consistent, not that it was invariant. Only a second, independently batched scoring path could reveal the difference.

**Table 2:** Dependence of M4 embeddings on inference batching, measured as the change in each query’s maximum cosine to TRAIN_REF. Same hash-verified checkpoint, same sequences, same device.

| comparison | mean $\vert \Delta\vert $ | max $\vert \Delta\vert $ | signed mean |
| --- | --- | --- | --- |
| sealed evaluator vs second scoring script (all ITS-core) | 0.0054 | 0.115 | — |
| batch 64 vs batch 256 (TEST_KNOWN ITS-core) | 0.0045 | 0.080 | — |
| batch 1 vs batch 64 (TEST_KNOWN ITS-core) | 0.0220 | 0.146 | −0.0110 |
| batch 1 vs batch 256 (TEST_KNOWN ITS-core) | 0.0226 | 0.151 | — |
| sealed evaluator vs batch 1 (TEST_KNOWN ITS-core) | 0.0226 | — | −0.0104 |

Batches of 64 and 256 are both heavily padded and differ little from each other; both differ from unpadded batch-1 inference by about five times as much. Between unpadded and padded scoring, 35.2% of TEST\_KNOWN ITS-core queries changed their nearest reference and 16.6% changed its genus. The magnitude of the shift correlates only weakly with sequence length (Spearman $\rho = -0.11$ ).

The artefact was largest where the sealed paper’s cross-view claim lay. Rescoring the M4 development matrix with exact-length inference lowered every cross-view cell and left the same-view cells almost unchanged (Table S5; data in `data/*_sealed`. `csv`). Spacer-to-core AUROC fell from 0.638 and 0.660 to 0.581 and 0.616, known-genus accuracy from 44.9% and 36.3% to 32.9% and 24.2%, and novel-family placement from 39.2% and 28.7% to 30.0% and 20.6%, whereas same-view AUROC moved by at most 0.015. This is consistent with padded normalisation adding a component shared across views, so that embeddings of different barcode regions looked more alike than they are. The artefact’s effect on AUROC was also not consistent in sign or size across partitions: correction lowered ITS2 AUROC by 0.023 on TEST, raised it by 0.003 on DEV, and moved the historical ITS2 cosine AUROC by only 0.001. It therefore cannot be corrected after the fact; only batch-invariant inference removes it.

Correcting the inference moved the TEST AUROCs from 0.754, 0.716 and 0.763 to 0.756, 0.699 and 0.740 (Table 5). The change was not symmetric noise: ITS-core rose slightly, while both spacers fell by about 0.02, so padding had been flattering the spacer results specifically.

### 3.2 The alignment baseline depends on its configuration

Two settings that are routinely left at their defaults each changed the alignment baseline substantially (Table 3; Figure 2). On the historical benchmark, the default termination heuristics returned a different best hit from an exhaustive search for 62.0% of novel-genus queries and 37.1% of known-genus ones, with mean identity gains of 2.93 and 1.30 percentage points. No hit ever worsened, as expected, since an exhaustive search cannot return a worse best hit than a truncated one. Only five of the 542 queries without a default-settings hit gained one, so the censored set reflects genuinely unalignable sequences rather than the rejection cap.

**Table 3:** How the alignment baseline depends on its configuration. (a) Default termination heuristics against an exhaustive search on the historical ITS2 bench-mark. (b) Query-coverage filtering on TEST_NOVEL: the ITS-core rows sweep the threshold and the spacer rows are the control, since ITS1 and ITS2 contain no conserved region.

| *(a) Default heuristics against exhaustive search, historical ITS2* |  |  |  |  |  |
| --- | --- | --- | --- | --- | --- |
| query set | with a hit | lower-identity hit | target changed | rescued | mean gain |
| LGO (novel genus) | 3,902 | 2,344 (60.1%) | 2,419 (62.0%) | 5 | +2.93 pp |
| LSO (known genus) | 3,243 | 1,108 (34.2%) | 1,203 (37.1%) | 0 | +1.30 pp |
| *(b) Novel-genus placement by query-coverage filter, TEST* |  |  |  |  |  |
| view | coverage filter | family | order | class |  |
| ITS-core | none | 53.7% | 61.5% | 64.9% |  |
|  | 0.5 | 73.1% | 94.5% | 96.3% |  |
|  | 0.7 | 81.2% | 96.6% | 98.4% |  |
|  | 0.8 | 84.6% | 97.0% | 98.6% |  |
|  | 0.9 | 85.8% | 97.0% | 98.7% |  |
| ITS1 | none | 80.4% | 94.4% | 96.5% |  |
|  | 0.8 | 81.6% | 94.4% | 96.5% |  |
| ITS2 | none | 76.2% | 95.3% | 97.6% |  |
|  | 0.8 | 80.0% | 95.6% | 97.5% |  |

**Figure 2:**
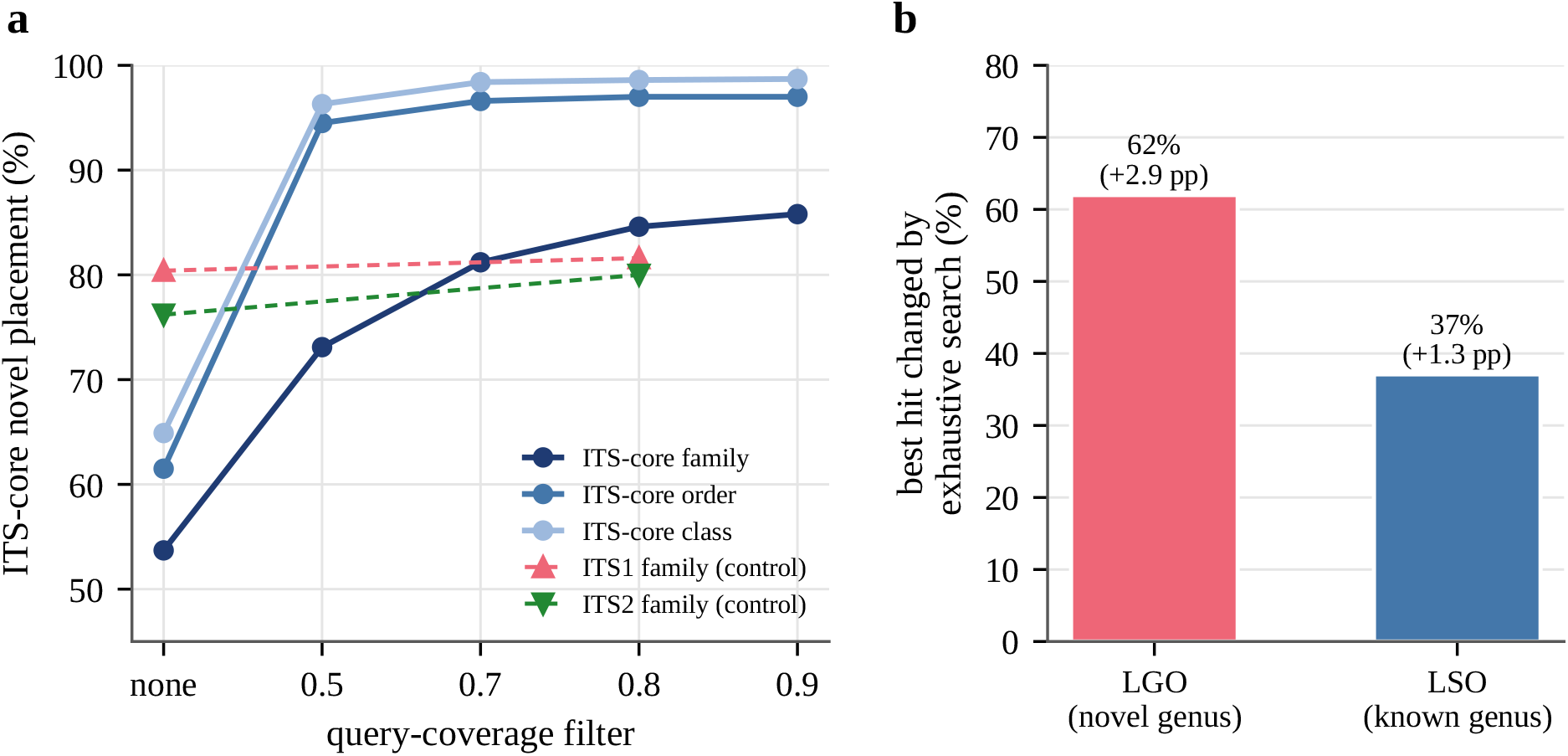
Configuration of the alignment baseline. (a) Best-hit placement of ITS-core novel genera as the query-coverage filter is tightened; dashed lines are the spacer controls, which contain no conserved region and barely move. (b) Share of historical-benchmark queries whose best hit changed when the default termination heuristics were replaced by an exhaustive search, with the mean identity gain.

The second setting was the coverage requirement. Without it, ITS-core identity placed novel genera in the correct family for only 53.7% and the correct class for 64.9%, which inverted the ordering expected from barcode length: ITS-core, which strictly contains both spacers, placed worse than either. Requiring 80% query coverage raised these to 84.6% and 98.7% and restored the expected ordering, while changing ITS1 and ITS2 family placement by only 1.2 and 3.8 points and their order and class placement not at all. The asymmetry identifies the cause. With no coverage requirement, a distant reference that aligns well over the conserved 5.8S can out-score a closer reference aligned across the full length; 44.5% of ITS-core best hits changed, with a mean identity change of −2.53 points, which is the signature of higher-identity partial alignments being replaced by correct full-length ones. The effect is insensitive to the exact threshold: order and class plateau from 0.7, and even 0.5 recovers most of it.

Strikingly, the coverage filter left identity’s novelty AUROC unchanged to three decimals at every view (Table 4). An evaluation of the alignment baseline by AUROC alone would therefore have reported a healthy baseline while one ITS-core novel query in three was being assigned to the wrong class.

**Table 4:** The coverage filter changes placement and leaves novelty ranking untouched. Identity on TEST under exhaustive search, with and without query coverage ≥ 0.8.

| view | configuration | AUROC | known genus NN | novel family |
| --- | --- | --- | --- | --- |
| ITS-core | exhaustive, no coverage filter | 0.7339 | 76.8% | 53.7% |
|  | exhaustive, query coverage 0.8 | 0.7338 | 83.4% | 84.6% |
| ITS1 | exhaustive, no coverage filter | 0.7016 | 80.4% | 80.4% |
|  | exhaustive, query coverage 0.8 | 0.6997 | 80.9% | 81.6% |
| ITS2 | exhaustive, no coverage filter | 0.7352 | 78.8% | 76.2% |
|  | exhaustive, query coverage 0.8 | 0.7349 | 80.8% | 80.0% |

### 3.3 A correctly configured alignment baseline matches or exceeds the encoder on the primary split

Scored on identical TEST queries and references, identity exceeded corrected M4 in known-genus nearest-neighbour accuracy at every view, 83.4%, 80.9% and 80.8% against 71.2%, 61.6% and 66.2%, and in novel-family placement at every view, 84.6%, 81.6% and 80.0% against 64.8%, 54.3% and 53.8% (Table 5; Figure 3). It also led at order and class. The sealed M4 values, shown alongside, do not change this at any view.

**Figure 3:**
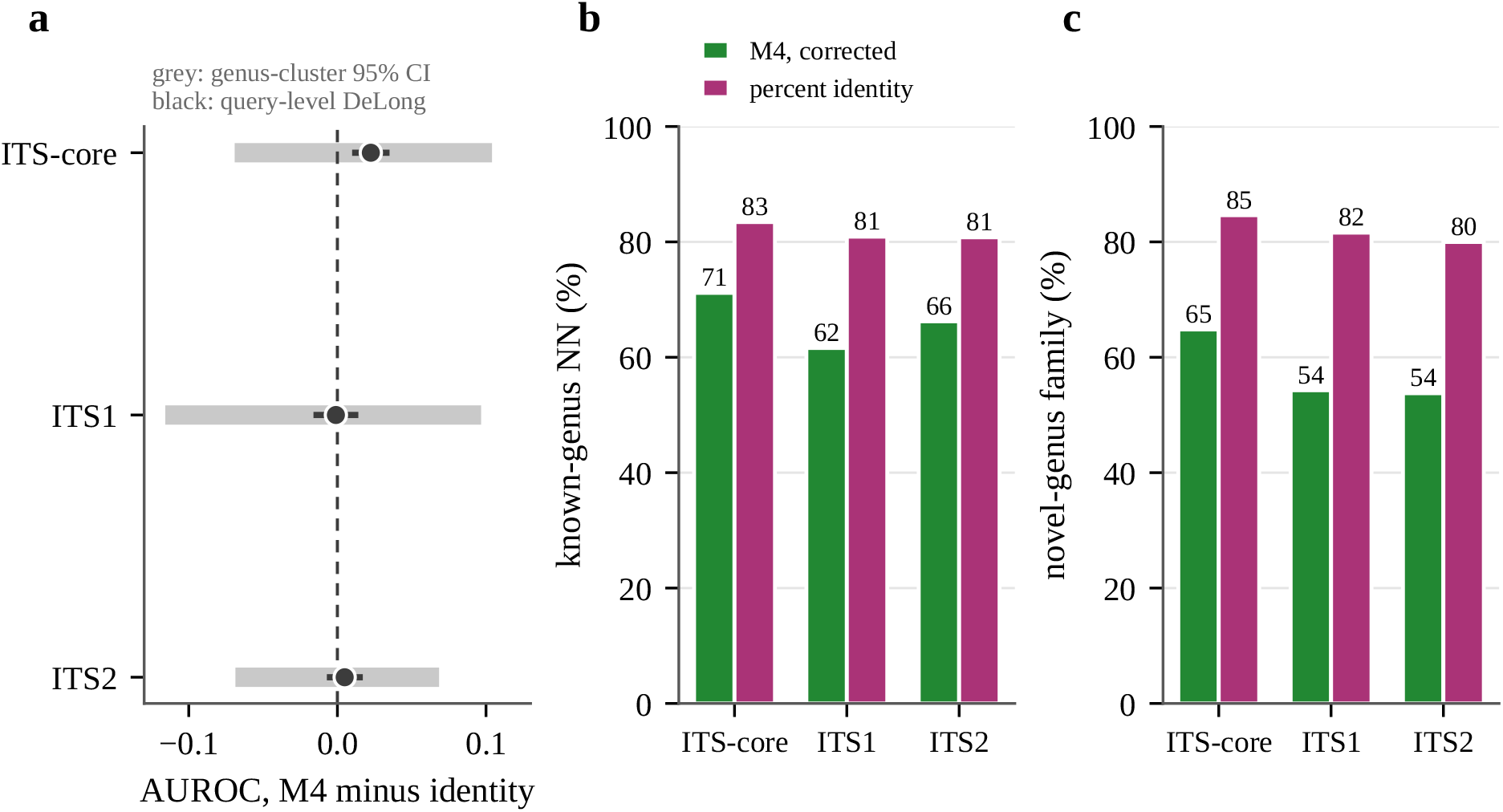
Corrected M4 against percent identity on identical TEST queries. (a) AUROC difference, M4 minus identity, with the query-level DeLong interval (black) and the genus-cluster bootstrap interval (grey). (b) Known-genus nearest-neighbour accuracy. (c) Novel-genus family placement.

**Table 5:** Primary TEST evaluation: M4 against a matched alignment baseline on identical queries and references. The sealed row is the locked evaluation as run, with padded inference; the corrected row recomputes the same checkpoint with exact-length batching. Identity is exhaustive VSEARCH with query coverage ≥ 0.8. Best value per view and column in bold.

| view | score | AUROC | known | novel-genus placement |  |  |
| --- | --- | --- | --- | --- | --- | --- |
|  |  |  | genus NN | family | order | class |
| ITS-core | M4, sealed (padded) | 0.754 | 71.1% | 65.0% | 92.2% | 97.3% |
|  | M4, corrected | **0.756** | 71.2% | 64.8% | 92.5% | 97.1% |
|  | percent identity | 0.734 | **83.4%** | **84.6%** | **97.0%** | **98.7%** |
| ITS1 | M4, sealed (padded) | **0.716** | 64.5% | 55.2% | 85.9% | 94.0% |
|  | M4, corrected | 0.699 | 61.6% | 54.3% | 84.4% | 93.3% |
|  | percent identity | 0.700 | **80.9%** | **81.6%** | **94.4%** | **96.5%** |
| ITS2 | M4, sealed (padded) | **0.763** | 65.3% | 52.0% | 81.1% | 92.9% |
|  | M4, corrected | 0.740 | 66.2% | 53.8% | 81.7% | 93.2% |
|  | percent identity | 0.735 | **80.8%** | **80.0%** | **95.6%** | **97.5%** |

$n$ known / novel: ITS-core 1,467/7,718; ITS1 1,477/7,702; ITS2 1,463/7,671. Identity censors queries with no hit at identity $\geq 0.5$ and coverage $\geq 0.8$ at 49.999 and counts them as placement failures: ITS-core 0 known, 0 novel; ITS1 26 known, 118 novel; ITS2 7 known, 78 novel.
Paired uncertainty for the AUROC and detection contrasts is in Table 6.

Novelty ranking was close. M4’s AUROC exceeded identity’s by +0.022 at ITS-core and +0.005 at ITS2 and trailed by -0.001 at ITS1 (Table 6). At the query level only the ITS-core difference was distinguishable from zero. The genus-cluster bootstrap widened every interval roughly fivefold, to -0.069 to +0.104, -0.116 to +0.097, and -0.069 to +0.068, so no view shows a demonstrated advantage for either score. The widening is itself informative: the novel population contains thousands of queries but only 183, 184 and 183 genera, and the between-genus variation that the query-level analysis ignores dominates the uncertainty. These intervals mean no demonstrated advantage, not equivalence: they admit material differences in either direction.

**Table 6:**
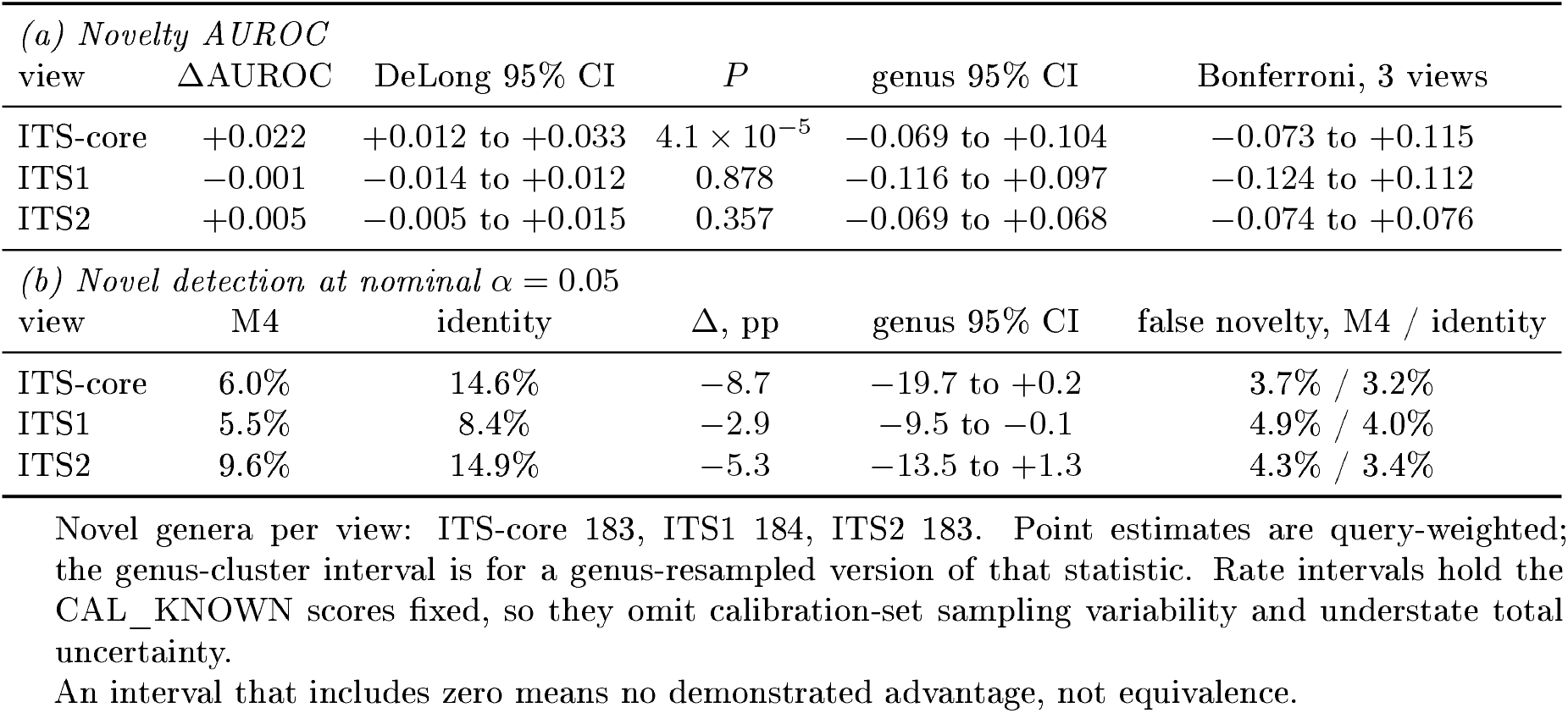
Paired uncertainty for corrected M4 against identity on TEST, M4 minus identity. The query-level DeLong interval treats queries as independent; the genus-cluster bootstrap resamples whole genera separately within the known and novel classes, applies identical weights to both scores, and uses 10,000 replicates.

### 3.4 Identity dominates at every deployable operating point

At *α* = 0.05, identity detected 14.6%, 8.4% and 15.0% of novel-genus queries against M4’s 6.0%, 5.5% and 9.6%, at observed false-novelty rates of 3.2%, 4.0% and 3.3% against 3.7%, 4.9% and 4.3% (Table 7; Figure 4). Identity therefore detected more and erred less simultaneously, at every view, and the pattern held at *α* = 0.10. At *α* = 0.01 both scores detected almost nothing, and ITS1 identity flagged no query at all. Under genus-cluster resampling the detection difference at *α* = 0.05 excluded zero at ITS1 and narrowly included it at ITS-core and ITS2.

**Table 7:** Conformal operating points on TEST for corrected M4 and identity, each calibrated on its own CAL_KNOWN scores by the same procedure. False novelty is the fraction of TEST_KNOWN flagged; detection the fraction of TEST_NOVEL flagged.

| view | $\alpha$ | M4, corrected | | percent identity | |
| --- | --- | --- | --- | --- | --- |
|  |  | false novelty | detection | false novelty | detection |
| ITS-core | 0.01 | 0.2% | **0.7%** | 0.5% | 0.3% |
|  | 0.05 | 3.7% | 6.0% | 3.2% | **14.6%** |
|  | 0.10 | 8.4% | 18.4% | 7.2% | **30.9%** |
|  | 0.20 | 17.6% | **45.3%** | 16.4% | 45.2% |
| ITS1 | 0.01 | 0.4% | **0.8%** | 0.0% | 0.0% |
|  | 0.05 | 4.9% | 5.5% | 4.0% | **8.4%** |
|  | 0.10 | 8.7% | 12.3% | 7.4% | **24.8%** |
|  | 0.20 | 18.1% | 30.6% | 16.9% | **45.4%** |
| ITS2 | 0.01 | 0.8% | **1.2%** | 0.5% | 1.1% |
|  | 0.05 | 4.3% | 9.6% | 3.3% | **15.0%** |
|  | 0.10 | 8.6% | 21.4% | 7.6% | **31.0%** |
|  | 0.20 | 17.5% | 41.1% | 17.2% | **44.9%** |

Higher detection per row in bold. At $\alpha \in \{0.05, 0.10\}$ identity detects more novel queries at a lower observed false-novelty rate at every view. At $\alpha = 0.01$ both scores detect almost nothing, and ITS1 identity flags no query at all.

**Figure 4:**
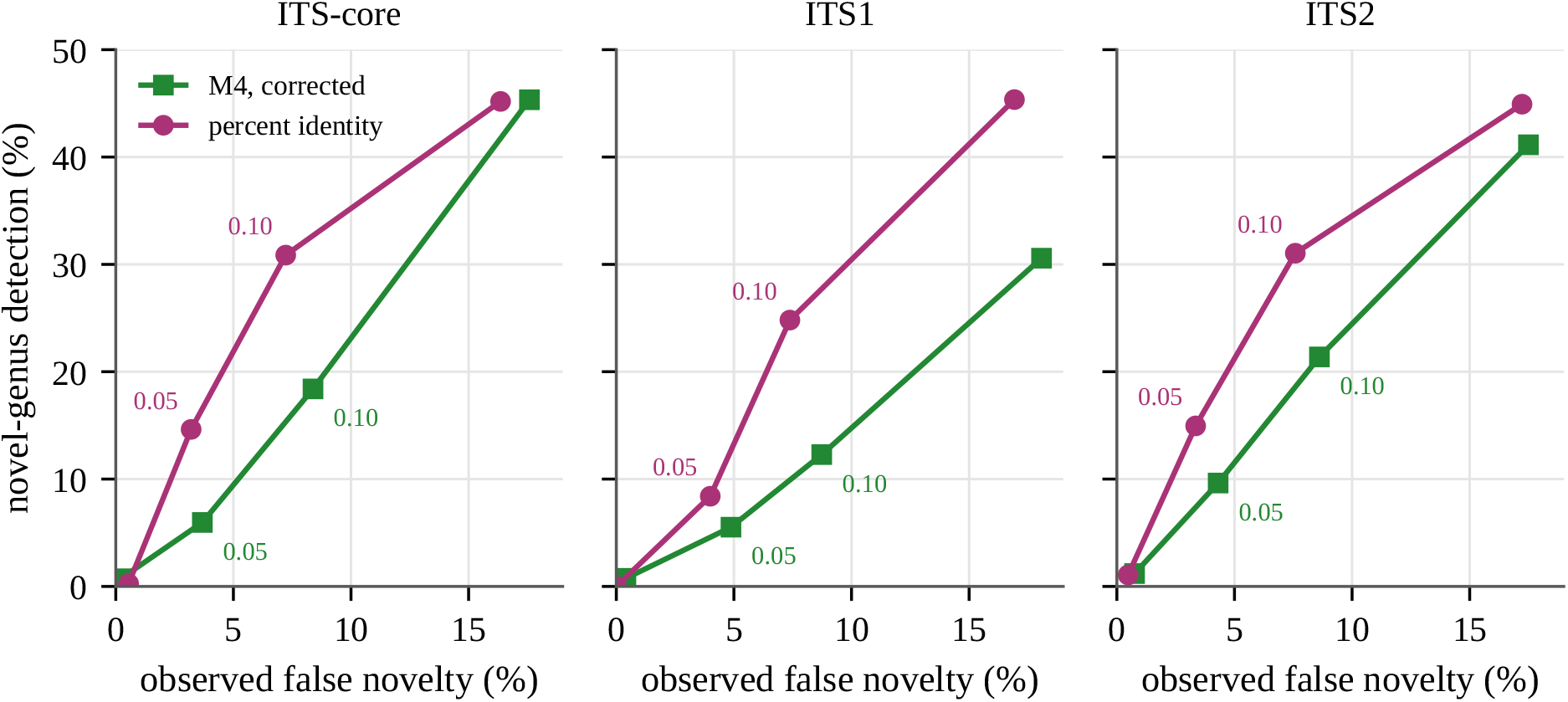
Conformal operating points on TEST. Each curve joins the nominal levels *α* = 0.01, 0.05, 0.10 and 0.20; the 0.05 and 0.10 points are labelled. A curve above and to the left of another dominates it. Identity dominates corrected M4 throughout the usable range at all three views.

This is the sharpest form of a point that applies well beyond this study. M4’s AUROC was comparable to identity’s, yet M4 was dominated in both coordinates at every operating point in the usable range. AUROC integrates over the whole curve, including high-error regions no user would choose; the decision a user makes is taken at a single stringent threshold. Novelty scores for barcoding should be compared at the operating points at which they would be deployed.

The novel genera flagged by the two methods at *α* = 0.05 overlapped only partly, with Jaccard indices of 0.17 to 0.23 and Spearman correlations between the scores of 0.61 to 0.72, so the two scores are not redundant. Their naive union was nonetheless worse than identity alone: flagging a query when either score did so detected 17.0%, 11.9% and 20.0% of novel queries at observed false-novelty rates of 5.9%, 7.9% and 6.6%, whereas identity alone, interpolated to the same error rates, detected about 25%, 26% and 27%. Any useful combination would require a rule fixed in advance and validated on a fresh holdout.

### 3.5 Cross-view retrieval is available to alignment through ITS-core

The sealed evaluation’s strongest cross-view results were retrieval of ITS1 or ITS2 queries against ITS-core references (Table S5). Because ITS-core contains both spacers, that comparison is homologous by construction, and an alignment can make it directly. On the same development queries and references, exhaustive identity reached known-genus accuracy of 79.8% and 78.5% against M4’s 32.9% and 24.2%, novel-family placement of 70.9% and 69.8% against 30.0% and 20.6%, and AUROC of 0.801 and 0.779 against 0.581 and 0.616 (Table 8). M4 was selected on these partitions, so these cells if anything flatter it, and the gaps are 47 to 54 points in genus accuracy and 41 to 49 in family placement. The one cross-view comparison alignment cannot make, direct ITS1-to-ITS2 retrieval, is also the one at which M4 performs at chance, with AUROCs of 0.490 and 0.471. The model’s cross-view ability is therefore core-anchored, and in the direction that matters it reproduces, less well, a retrieval alignment performs without training.

**Table 8:**
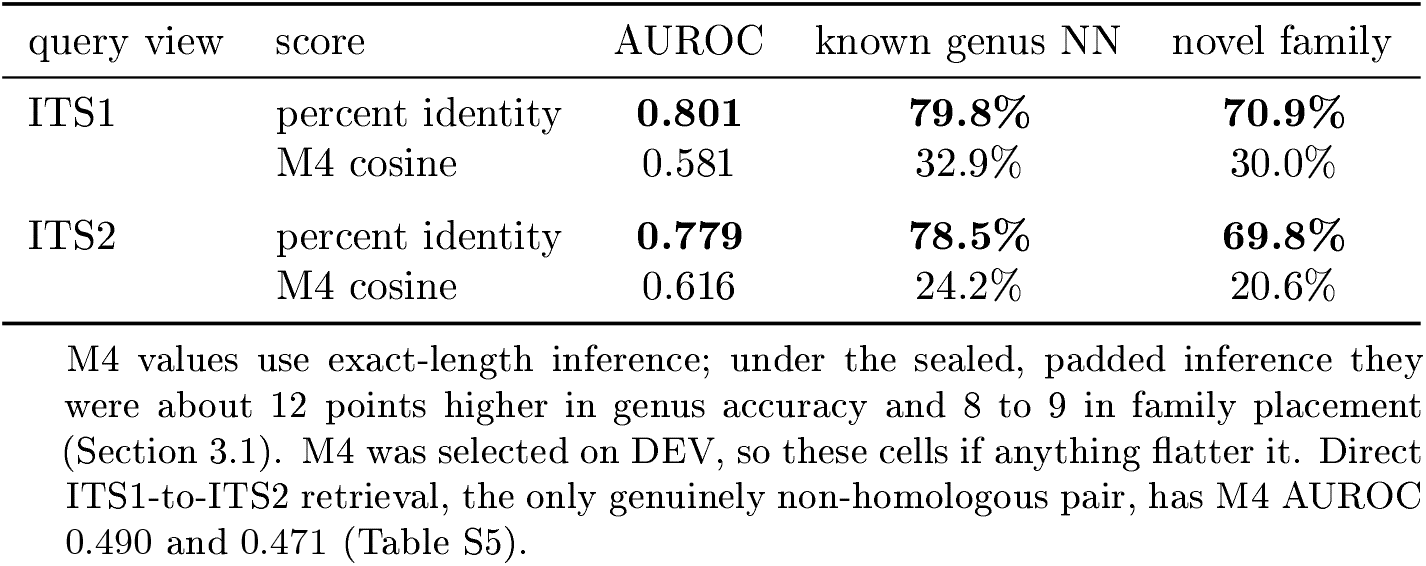
Cross-view retrieval through the ITS-core references on DEV. Spacer queries searched against ITS-core references, by exhaustive VSEARCH (query coverage ≥ 0.8) and by M4 cosine. Because ITS-core contains both spacers, this is a homologous comparison that alignment can make directly.

### 3.6 The development-to-TEST gap is partition composition, not selection

M4’s development AUROCs lie above its corrected TEST values at every view, by +0.021, +0.041 and +0.014 (Table 9). A learned score cannot separate selection optimism from a difference between partitions on its own, so we scored identity on both. Identity is parameter-free and nothing was selected on it, so its development-to-TEST gap estimates partition composition alone. That gap was larger than M4’s at every view, +0.075, +0.107 and +0.044, leaving no positive residual to attribute to selection. Family placement agreed: both scores were 7 to 14 points lower on DEV_NOVEL than on TEST_NOVEL. DEV_NOVEL is simply the easier partition for novelty ranking and the harder one for placement. Every value in this comparison uses exact-length inference or exhaustive search. A development-above-test gap is often read as selection optimism; here a parameter-free control shows that reading to be wrong on the data that produced it.

**Table 9:**
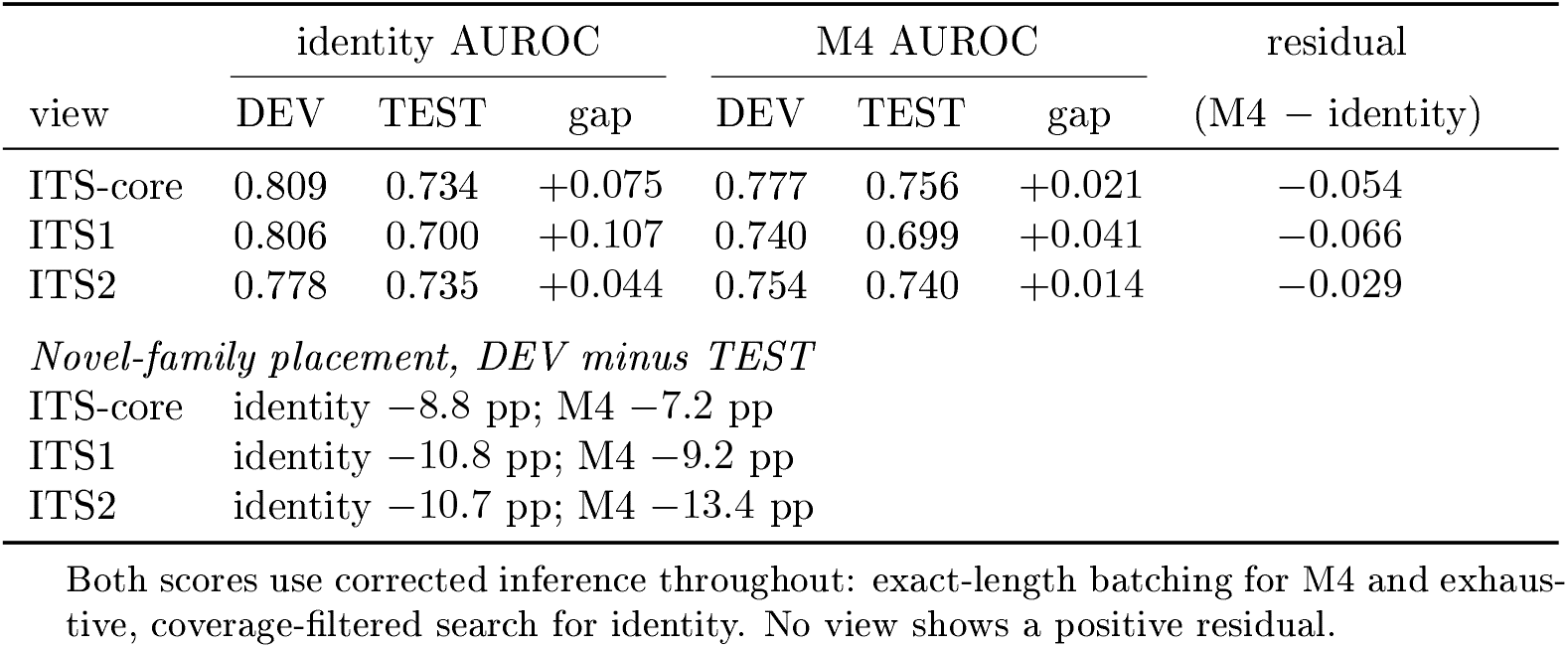
Development-to-TEST movement separated into partition composition and selection. Identity is parameter-free and nothing was selected on it, so its gap estimates composition alone. Same-view retrieval throughout.

| view | identity AUROC |  |  | M4 AUROC |  |  | residual<br>(M4 – identity) |
| --- | --- | --- | --- | --- | --- | --- | --- |
|  | DEV | TEST | gap | DEV | TEST | gap |  |
| ITS-core | 0.809 | 0.734 | +0.075 | 0.777 | 0.756 | +0.021 | −0.054 |
| ITS1 | 0.806 | 0.700 | +0.107 | 0.740 | 0.699 | +0.041 | −0.066 |
| ITS2 | 0.778 | 0.735 | +0.044 | 0.754 | 0.740 | +0.014 | −0.029 |
| *Novel-family placement, DEV minus TEST* |  |  |  |  |  |  |  |
| ITS-core | identity −8.8 pp; M4 −7.2 pp |  |  |  |  |  |  |
| ITS1 | identity −10.8 pp; M4 −9.2 pp |  |  |  |  |  |  |
| ITS2 | identity −10.7 pp; M4 −13.4 pp |  |  |  |  |  |  |

Both scores use corrected inference throughout: exact-length batching for M4 and exhaustive, coverage-filtered search for identity. No view shows a positive residual.

### 3.7 A historical benchmark was contaminated by training data

The secondary benchmark reuses the leave-one-genus-out and leave-one-species-out workflow of our companion study, built from the same UNITE release. Before scoring it we audited whether its held-out taxa had appeared in M4 training. 3,166 of the 4,444 novel-genus queries (71%) and 458 of the 538 held-out genera (85%) were present in the original TRAIN_REF (Table S2; Figure S1). Scoring the frozen checkpoint on this benchmark would have measured recall of training data under the name of novelty detection. We therefore retrained the fixed recipe on a TRAIN_REF with every leaked taxon removed. The mechanism is general: any leave-one-taxon-out benchmark derived from the same release as a model’s training data will leak unless exclusions are enforced before training rather than at evaluation. On the leakage-safe benchmark, identity again outperformed the retrained model, and here the difference is statistically clear. AUROC was 0.768 against 0.703, a difference of +0.065 whose genus-clustered interval, resampling 538 novel and 1,387 known genera, ran from +0.046 to +0.085 (Table S6; Figure S4). At *α* = 0.05 identity detected 28.2% of novel-genus queries against 11.2%, with 872 queries flagged by identity alone and 117 by cosine alone, and it recovered the correct family for 52.6% of them against 28.2%. About half of identity’s detections were queries with no alignable reference at all. Two features of the benchmark plausibly make it harsher for the model than the primary split: the retrained checkpoint was fitted to 11% fewer records and 483 fewer genera, and its known class consists of held-out species rather than known genera.

## 4 Discussion

Every comparison between the learned embedding and a correctly configured alignment baseline either favoured alignment or showed no demonstrated difference. Alignment placed novel genera more accurately at every view and rank, identified known genera more accurately, detected more novel genera at a lower error rate at every usable threshold, and performed the cross-view retrievals credited to the embedding far better. The embedding retained a novelty AUROC close to identity’s, which the operating-point analysis showed to be the least decision-relevant of the metrics examined.

The reason is not mysterious. For homologous queries and references, alignment compares sequences at full nucleotide resolution. The encoder compresses the same information into 256 dimensions while being asked to satisfy locus invariance, genus discrimination and hierarchical structure at once. Where the compression could in principle add something alignment lacks, in a shared space for non-homologous barcode views, the model performed near chance, because the cross-view structure it did learn was anchored on the ITS-core region that alignment can use directly.

The more transferable result concerns the evaluation. The sealed evaluation was carefully designed, with a genus-separated split, a sealed test set, a hash check and preregistered integration targets, and it still supported a conclusion that did not survive. Each correction that changed it is inexpensive:

1. Score a non-learned baseline on the same split, not only weaker versions of the model.
2. Configure that baseline as a user would for best-hit assignment: exhaustive search, and a coverage requirement when the barcode contains a conserved region. Report the configuration.
3. Compare novelty scores at deployable operating points, not only by AUROC, which here was blind both to a broken baseline and to a dominated model.
4. Compute uncertainty at the level of the unit that is actually independent, here the genus.
5. Test whether embeddings are invariant to inference batching by scoring the same queries through two independently batched paths.
6. Use a parameter-free score as the control for selection optimism.
7. Audit any benchmark for training-set leakage before scoring it.

None of these requires a new method. Each changed a conclusion here, and several would have been invisible without the others: the coverage artefact was invisible to AUROC, and the padding effect was invisible to exact reproduction.

### 4.1 Relation to prior work

Orsholm et al. compared classifiers for unknown barcodes across animal COI and fungal ITS and found composition-based classifiers strongest for fungal ITS, with alignment-threshold methods over-assigning queries to novel branches [25]. Our results are consistent with theirs on the importance of baseline configuration and complementary in scope: they compared classification rules across many methods, whereas we examined how evaluation choices decide a single learned-versus-alignment comparison. Zero-shot and hierarchical Bayesian models for insect barcodes [2, 3] and out-of-distribution detection for insect barcoding [5] share our motivation, and the last reaches a compatible conclusion, that learned scores do not automatically dominate distance baselines. Benchmark design has shaped earlier conclusions in the same way. Edgar’s cross-validation-by-identity benchmark for 16S and fungal ITS found that newer classification algorithms did not improve on the RDP classifier or SINTAX [24], and the SINTAX evaluation showed most methods assigning known names to novel taxa [23]. Our results extend that line from classification rules to learned representations. Established probabilistic classifiers with explicit novelty handling, including PROTAX-Fungi [21] and BayesANT [26], and composition classifiers such as SINTAX [23], are the natural next baselines for the protocol described here.

### 4.2 Limitations

All experiments use one UNITE release. The strongest test of generalisation, training on an earlier release and evaluating taxa that first appear in a later one, remains to be done, and would also address whether learned barcode embeddings largely memorise their training set.

The leakage-safe retrain was not re-evaluated on DEV, so the cost of the exclusion mask to model quality is unmeasured, and the historical comparison may understate what an unmasked model would achieve on genuinely novel taxa.

The ladder was run at a single seed, so small differences between adjacent rungs are not separable from run-to-run variation. The genus-cluster bootstrap assumes genera are independent within each class and ignores family-level dependence, and its rate intervals hold the calibration set fixed, so they understate total uncertainty. The coverage threshold was chosen after unfiltered TEST placement had been seen; the sweep shows the conclusion does not depend on the value, and the filter improves the baseline rather than the model under study. We reported all three barcode views with equal weight and did not select among them after seeing results.

## 5 Conclusions

A purpose-trained fungal ITS embedding did not outperform correctly configured alignment on any comparison we could make, and fell well short of it on higher-rank placement, genus identification, deployable novelty detection and cross-view retrieval. The comparison was decided by evaluation choices: how the alignment baseline was configured, whether the embedding was invariant to batching, whether uncertainty was computed at the genus level, whether a parameter-free control was used for selection, and whether the benchmark had leaked into training. These checks are general and inexpensive, and we suggest that learned barcode methods be reported against them.

## Data and code availability

Code, the deviation log with its commit history, and the environment files are de-posited at https://github.com/ayobi/openvorld-its-paper and archived on Zenodo (doi:10.5281/zenodo.22940788). Every table, figure and quoted number in this manuscript is re-generated from the CSV files in `manuscript/data` by `manuscript/analysis/make_tables.py, make_figures. Py` and `make_primary.py;` the historical-benchmark CSV is itself rebuilt from the per-query score table and the raw best-hit tables by `analysis/rebuild_historical_benchmark.py`, and the per-query table by `scripts/build_historical_per_query.py.` Model checkpoints, the split table, and the per-query predictions and search hits are deposited separately on Zenodo (doi:10.5281/zenodo.22940616). The sequence source is the UNITE general FASTA release for Fungi of 19 February 2025 [17].

## Author contributions

Aaron O’Brien: conceptualization, methodology, software, formal analysis, investigation, data curation, visualization, writing (original draft), writing (review and editing) . Auguste Gardette: conceptualization, methodology, validation, writing (review and editing) . A.G. contributed to the conception of the study and proposed its framing as an evaluation protocol. His review identified the absence of a non-learned baseline on the primary split, the homologous confounder in cross-view retrieval, and the termination behaviour of the default VSEARCH search, each of which led to an analysis reported here. Both authors read and approved the submitted version.

### Acknowledgements

We thank the UNITE community for curating and releasing the reference data on which every experiment here depends.

## Use of generative AI

Anthropic’s Claude and GitHub Copilot were used in preparing this work: to draft and revise manuscript text, to write and debug the analysis and figure scripts deposited with it, and to check bibliographic details. All analyses were run by the authors, all reported values were verified against the primary output, and the authors are responsible for the content of the manuscript.

### A Reproducibility, frozen artefacts and supplementary results

The primary frozen M4 checkpoint SHA-256 is

`e4cf8bfed197b2b15758a93798c025a2a482e5ad81c5965b376bcf4bb237671f.`

The split TSV SHA-256 is

`27041e67b8db254c31a8fcb4001b35c07486786777d6ce09a21a86c4997cb411.`

The locked final evaluator embedded TRAIN_REF, CAL_KNOWN, TEST_KNOWN and TEST_NOVEL but did not embed or score DEV. The historical-safe retraining used the selected M4 recipe unchanged and skipped DEV, CAL and TEST evaluation entirely. The leakage-safe checkpoint scored in Table S6 has SHA-256

`99b91448b0401acbc6d007156df08dc85420036bffdc80ef54746c11280de16a.`

Each corrected evaluation records its evaluator and model-source hashes and its inference policy in its output metadata, and every one of those hashes matches the corresponding file deposited in the repository.

Neural training was run with PyTorch 2.14.0+cu130 on an NVIDIA GeForce RTX 5070 Ti with CUDA 13.0; ITS extraction used ITSx 1.1.3. The training environment is deposited as `environment.` lock. The corrected evaluation and all analyses ran on Python 3.13.12 (conda-forge), PyTorch 2.14.0 with CUDA 13.0, NumPy 2.5.3, pandas 3.0.6 and SciPy 1.18.1, on the same GPU (driver 595.84), with VSEARCH v2.31.0; the full package list is deposited as `environment/requirements.txt.` Hashes prove that an artefact did not change; the environment files are what make it rebuildable.

The run logs deposited with the code date each fit of the ladder: M0 through M3 completed between 12:34 and 13:48 on 11 September 2026 and M4 began at 13:57, so the sequence on which the integration targets depend is checkable from the artefacts rather than only asserted. The threshold values themselves were fixed in working notes in that interval and are not independently timestamped (§2.2). Paired uncertainty for the historical benchmark is computed by `analysis /paired_uncertainty.Py`, which reports a clustered paired bootstrap interval for the AUROC difference, the DeLong test for the same contrast, and McNemar’s test on the conformal flag sets at each *α*.

**Table S1:** Sensitivity of the hierarchical objective to its weight. The reduced *λ*_tax_ = 0.25 run was performed once after the initially specified *λ*_tax_ = 1.0. run and was not followed by a weight sweep.

| $\lambda_{\text{tax}}$ | view | AUROC | novel family | novel order |
| --- | --- | --- | --- | --- |
| 1.00 | ITS-core | 0.752 | 35.8% | 60.8% |
|  | ITS1 | 0.740 | 32.7% | 56.9% |
|  | ITS2 | 0.740 | 31.5% | 56.9% |
| 0.25 | ITS-core | 0.748 | 33.6% | 58.7% |
|  | ITS1 | 0.736 | 28.8% | 52.2% |
|  | ITS2 | 0.723 | 29.7% | 53.8% |

Reducing the weight to one quarter lowered AUROC and every placement metric at every view. The sensitivity run therefore did not motivate further DEV-set weight tuning.

**Table S2:** Historical-benchmark leakage audit and exclusion mask. The share column expresses each count as a percentage of the corresponding held-out set.

| item | count | share of held-out set |
| --- | --- | --- |
| *Training records* |  |  |
| original TRAIN_REF | 46,453 |  |
| leakage-safe TRAIN_REF | 41,344 |  |
| excluded by the mask | 5,109 |  |
| *Leave-one-genus-out (LGO) queries* |  |  |
| queries | 4,444 |  |
| query genera | 538 |  |
| exact query IDs in original TRAIN_REF | 3,166 | 71.2% |
| query genera in original TRAIN_REF | 458 | 85.1% |
| query genera after the mask | 0 | 0.0% |
| exact query IDs after the mask | 0 | 0.0% |
| *Leave-one-species-out (LSO) queries* |  |  |
| queries | 3,264 |  |
| query species | 2,541 |  |
| exact query IDs in original TRAIN_REF | 2,107 | 64.6% |
| query species in original TRAIN_REF | 1,694 | 66.7% |
| query species after the mask | 0 | 0.0% |
| query genera still represented after the mask | 1,887 |  |

The LSO design deliberately keeps the query’s genus in the reference collection, so the last row is a property of the design rather than residual leakage. What the mask had to remove was the query species itself.

**Figure S1:**
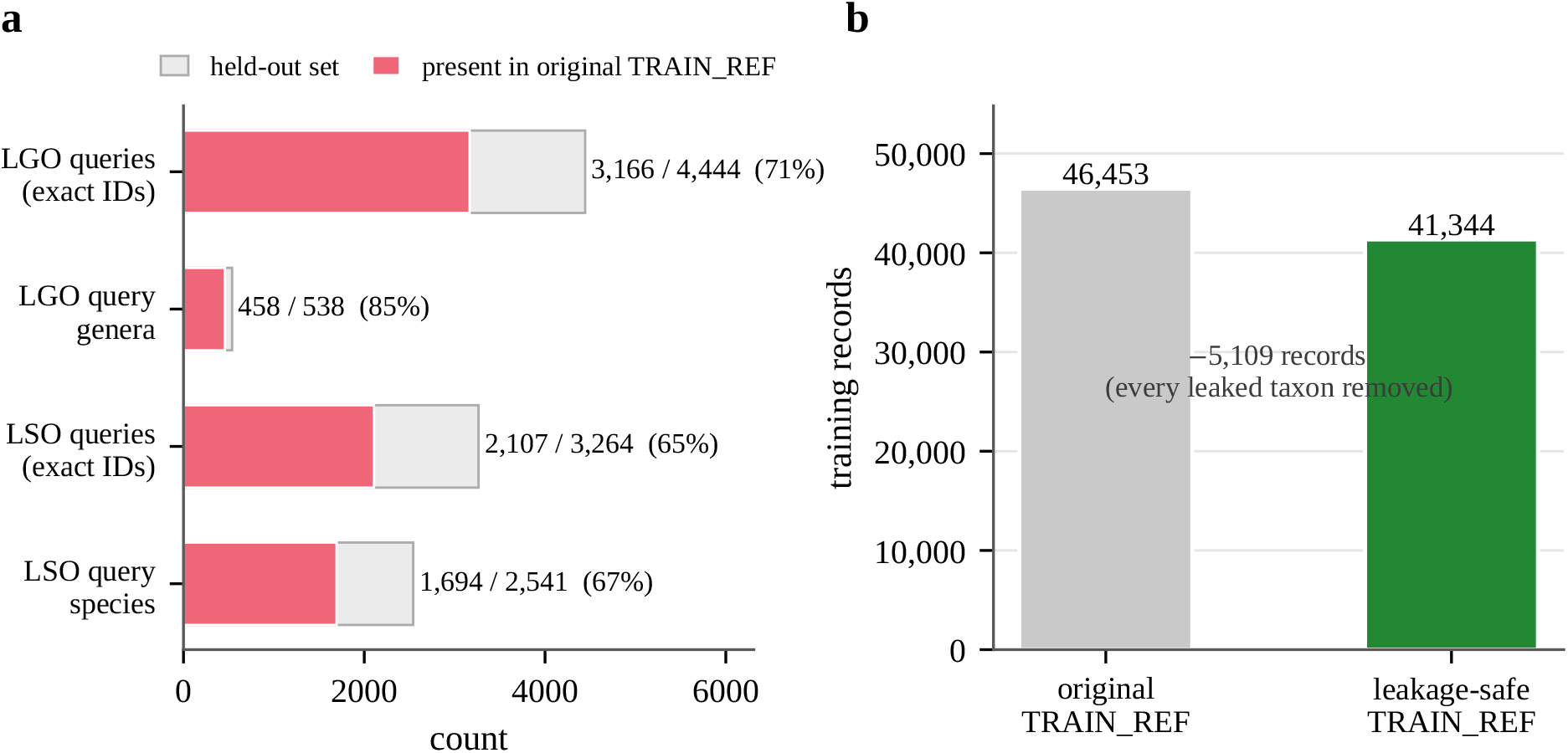
Why the frozen checkpoint could not be scored on the historical benchmark. (a) Most nominally held-out historical queries and query taxa were already present in the original TRAIN_REF. (b) Removing every one of them cost 5,109 training records and brought residual leakage to zero, after which the fixed M4 recipe was retrained.

## B Development ladder and training objectives

### B.1 Training objectives

Let a training batch contain *B* source records. Each record contributes an ITS-core embedding 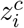 and one randomly selected available spacer embedding 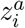 (ITS1 or ITS2). Let *w*_*g*_ denote a learned, L2-normalized proxy for genus *g*.

#### Genus proxy (all models)

The genus head is a cosine-softmax classifier evaluated on ITS-core only:

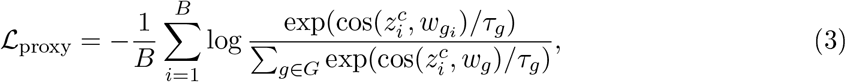

with *τ*_*g*_ = 0.07.

#### Cross-view invariance (Ml-M4)

The implementation uses a symmetric supervised-contrastive objective over all 2*B* views in the batch. If *π*(*a*) denotes the other view from the same source record as anchor *a*, then

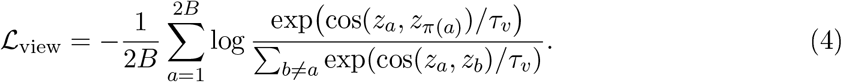

Thus both core→spacer and spacer→core directions are anchors, and every non-self view in the minibatch enters the denominator. We used *τ*_*v*_ = 0.10.

#### Hierarchical laxonomic geometry (M2, M4)

This term acts on pairwise ITS-core embeddings within the minibatch, not on genus proxies. For records *i* and *j*, let *h*_*ij*_ ∈ {0,…,6} be the number of consecutive ranks shared when their lineages are traversed from phylum toward species until the first mismatch, and define *d*_*ij*_ = 6 − *h*_*ij*_. For *j* ≠ *i*, the target and embedding-neighbour distributions are

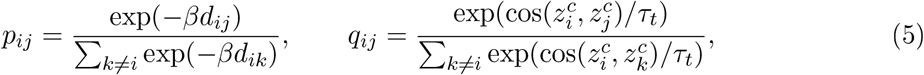

and

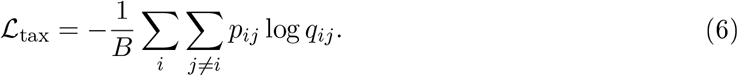

We used *τ*_*t*_ = 0.10 and *β* = 0.70.

#### Leave-one-genus family episodes (M3, M4)

Family prototypes are constructed from all learned genus proxies. Writing 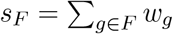 and *c*_*F*_ = norm(*s*_*F*_), an eligible query from genus *g*_*i*_ and family *F*_*i*_ replaces only its true-family prototype with

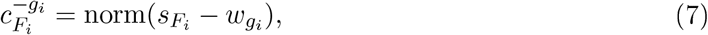

which explicitly removes the query genus. Families represented by only one genus are skipped. The query is classified over all represented families, using 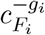 for the target logit and ordinary *c*_*F*_ for every non-target family:

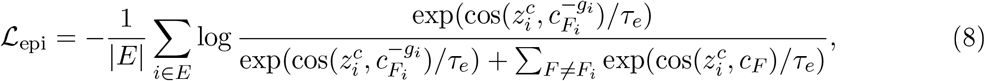

where *E* is the set of eligible queries and *τ*_*e*_ = 0.10.

The total objective is *L* = *L*_proxy_ + *λ*_view_*L*_view_ + *λ*_tax_*L*_tax_ + *λ*_epi_*L*_epi_, with the weights in Table S3.

**Table S3:** Controlled model ladder. Each step adds exactly one scientific ingredient to the step before it; M4 is the combination that was frozen. M0 was rerun at batch size 96 so that every comparison shown here shares the same source-record batch size and eight-epoch update budget.

| model | training objective added | $\lambda_{\text{view}}$ | $\lambda_{\text{tax}}$ | $\lambda_{\text{epi}}$ |
| --- | --- | --- | --- | --- |
| M0 | genus proxy on ITS-core only (Eq. 3) | — | — | — |
| M1 | + cross-view invariance, core↔ITS1/ITS2 (Eq. 4) | 1.0 | — | — |
| M2 | + hierarchical taxonomic geometry (Eq. 6) | 1.0 | 1.0 | — |
| M3 | + leave-one-genus family episodes (Eq. 8) | 1.0 | — | 0.5 |
| M4 | + both structured objectives | 1.0 | 1.0 | 0.5 |

All models share the same convolutional encoder and 256-dimensional L2-normalized output. View, taxonomy and episode temperatures were all 0.10; the hierarchical decay was $\beta = 0.70$ ; seed 17; eight epochs; batch size 96.
M2 and M3 are both built on M1, so the M2/M3 contrast isolates hierarchical geometry against pseudo-novel episodes at equal view supervision.

All controlled runs used eight epochs, batch size 96 and seed 17. Training records were sampled with replacement using inverse-genus-frequency weights. Optimization used AdamW (learning rate 3 × 10^−4^, weight decay 10^−4^) and cosine annealing to zero over eight epochs. Mixed-precision FP16 training was enabled on an NVIDIA GeForce RTX 5070 Ti. The primary M4 loader contained 46,150 paired records from 4,231 genera and 897 families, one fewer than the 46,151 ITS-core records in Table 1 because a single record carries an annotated core span but no annotated spacer and so forms no view pair; 3,872 genera belonged to families with at least two represented genera and were eligible for the episodic term. M1—M4 used *λ*_view_ = 1.0; M2 and M4 used *λ*_tax_ = 1.0; M3 and M4 used *λ*_epi_ = 0.5. An initial M0 run at batch size 256 exposed an update-budget confound, since a larger batch at fixed epochs means fewer optimizer steps. M0 was therefore rerun at batch size 96 and only that batch-matched run enters Table S4.

### B.2 Development metrics

Known queries are scored by nearest-neighbour genus accuracy. For DEV_NOVEL, where the true genus is absent by construction, we report whether the nearest reference falls in the true family, order or class. Novelty AUROC treats held-out genera as the positive class and *A*(*q*) as the score. To test locus invariance directly, each query view was also searched against each of the three reference views, producing a 3 × 3 retrieval matrix.

### B.3 Corrected development ladder

All values in this subsection use exact-length inference on the saved checkpoints; the sealed, padded values are deposited as `data/*s_sealed.Csv`. Correction left the ordering M0 *<* M1 *<* M4 intact at every view. It reversed the sign of the M3-versus-M1 AUROC diflerence at ITS2, from −0.008 to +0.013, so the trade-ofl between pseudo-novel episodes and novelty discrimination survives only at ITS-core, where it is small (−0.005). The M4 cross-view cells fell substantially (Section 3.1).

**Table S4:** Development-set ablation, same-view retrieval. Novelty AUROC treats held-out genera as the positive class and −*S*(*q*) as the score. The placement columns give the proportion of novel-genus queries whose nearest TRAIN_REF neighbour belongs to the true family, order and class. Best value per column in bold.

| model | view | AUROC | known | novel-genus placement |  |  |
| --- | --- | --- | --- | --- | --- | --- |
|  |  |  | genus NN | family | order | class |
| M0 | ITS-core | 0.681 | 49.1% | 32.5% | 57.1% | 82.9% |
|  | ITS1 | 0.676 | 45.6% | 28.2% | 50.8% | 75.1% |
|  | ITS2 | 0.684 | 47.4% | 24.3% | 48.0% | 76.5% |
| M1 | ITS-core | 0.748 | 59.2% | 34.0% | 56.8% | 85.7% |
|  | ITS1 | 0.732 | 54.6% | 29.2% | 51.0% | 80.9% |
|  | ITS2 | 0.710 | 51.7% | 30.0% | 54.0% | 83.4% |
| M2 | ITS-core | 0.752 | 63.3% | 35.8% | 60.8% | 89.3% |
|  | ITS1 | 0.740 | 56.9% | 32.7% | 56.9% | 84.2% |
|  | ITS2 | 0.740 | 57.1% | 31.5% | 56.9% | 83.8% |
| M3 | ITS-core | 0.743 | 64.8% | 52.4% | 78.7% | 93.9% |
|  | ITS1 | 0.733 | 57.6% | 38.7% | 63.4% | **86.9%** |
|  | ITS2 | 0.723 | 56.6% | 38.8% | 66.5% | 87.1% |
| M4 | ITS-core | **0.777** | **69.8%** | **57.6%** | **80.5%** | **94.7%** |
|  | ITS1 | **0.740** | **61.8%** | **45.1%** | **67.1%** | 86.1% |
|  | ITS2 | **0.754** | **61.7%** | **40.3%** | **66.9%** | **89.0%** |

M2 denotes the primary $\lambda_{\text{tax}} = 1.0$ run; the reduced-weight sensitivity run is Table S1.
Single run per configuration (seed 17). Between-seed variability was not estimated; differences of approximately 0.01 AUROC between adjacent rungs should therefore be interpreted cautiously.

## C Historical benchmark

Identity from the exhaustive, coverage-filtered search and cosine from exact-length inference on the leakage-safe checkpoint. The sealed values are deposited as `data/historical_benchmark_sealed. Csv`.

**Figure S2:**
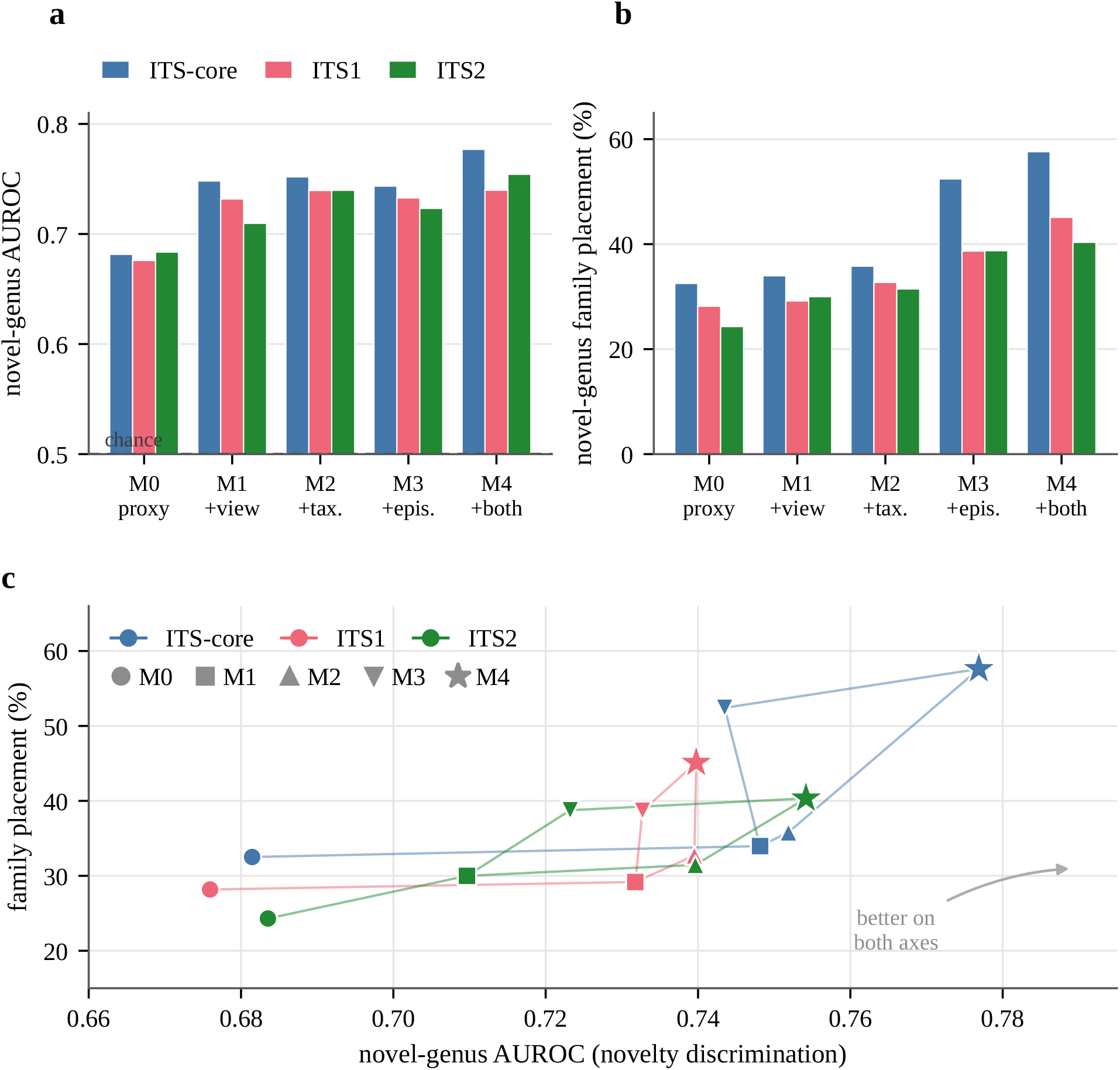
Development ladder, exact-length inference. (a) Novel-genus AUROC, axis truncated at chance. (b) Novel-genus family placement. (c) The two plotted against each other. Colour is the barcode view and marker shape the model.

**Table S5:** M4 development retrieval matrix. Every query view was searched against every reference view. Diagonal cells are same-view retrieval; off-diagonal cells measure how far the shared space transfers. Same-view cells in bold.

| metric | query view | reference view |  |  |
| --- | --- | --- | --- | --- |
|  |  | ITS-core | ITS1 | ITS2 |
| novelty AUROC | ITS-core | **0.777** | 0.598 | 0.615 |
|  | ITS1 | 0.581 | **0.740** | 0.490 |
|  | ITS2 | 0.616 | 0.471 | **0.754** |
| known-genus NN accuracy | ITS-core | **69.8%** | 37.0% | 27.5% |
|  | ITS1 | 32.9% | **61.8%** | 2.8% |
|  | ITS2 | 24.2% | 3.3% | **61.7%** |
| novel-genus family NN | ITS-core | **57.6%** | 36.3% | 30.1% |
|  | ITS1 | 30.0% | **45.1%** | 9.0% |
|  | ITS2 | 20.6% | 6.3% | **40.3%** |

ITS-core is the strongest bridge to both spacers. Direct ITS1↔ITS2 retrieval remains near chance in AUROC and below 7% in known-genus accuracy, consistent with incomplete alignment of the two spacer-specific representations.
Chance for a nearest-neighbour genus assignment over the 4,231 represented genera is approximately 0.02%, so the spacer-to-spacer cells are weak rather than empty.

**Figure S3:**
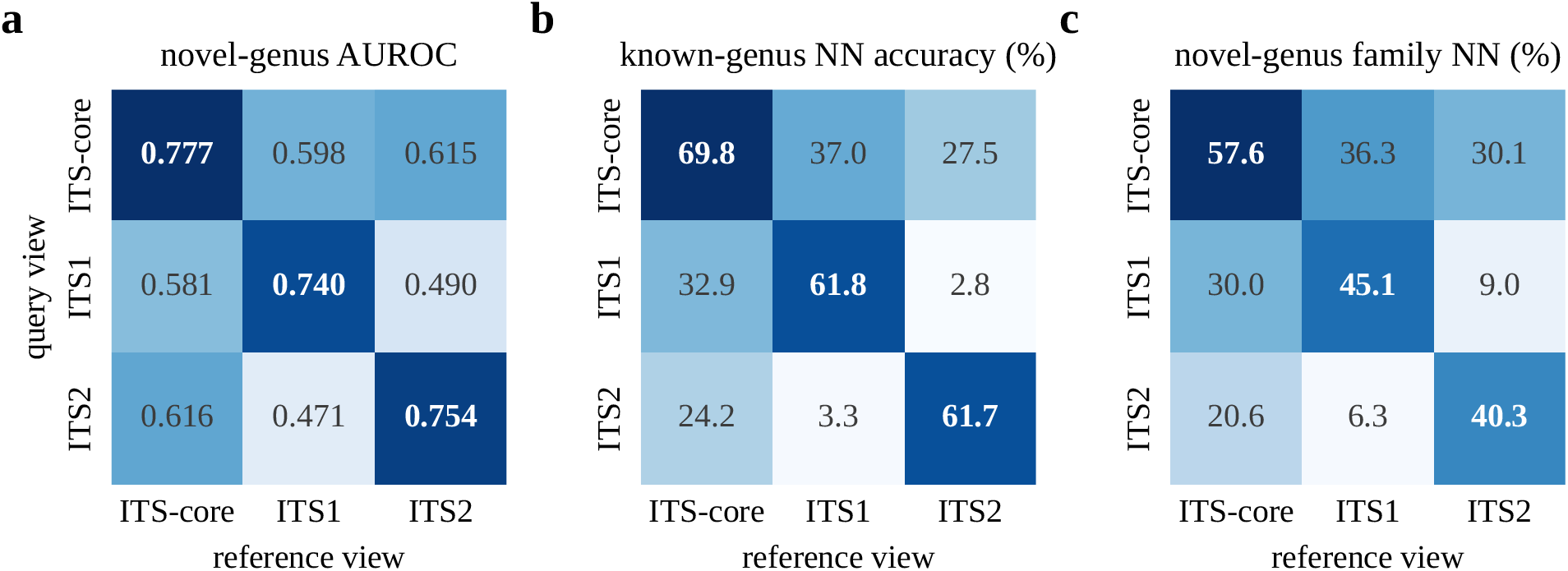
M4 development retrieval for every query and reference view, exact-length inference: (a) novel-genus AUROC, (b) known-genus nearest-neighbour accuracy, (c) novel-genus family placement.

**Table S6:** Leakage-controlled historical ITS2 benchmark. Percent identity and leakage-safe M4 cosine are scored on identical queries, identical reference collections and an identical conformal procedure, so the comparison is paired throughout. Higher is better except for false novelty, where the target is the nominal *α*.

| metric | percent identity | M4 cosine | difference |
| --- | --- | --- | --- |
| AUROC | **0.768** | 0.703 | +0.065 |
| novel detected, $\alpha = 0.01$ | **14.9%** | 2.4% | +12.5 pp |
| novel detected, $\alpha = 0.05$ | **28.1%** | 11.2% | +17.0 pp |
| novel detected, $\alpha = 0.10$ | **38.5%** | 20.9% | +17.6 pp |
| novel detected, $\alpha = 0.20$ | **54.0%** | 41.0% | +13.0 pp |
| false novelty, $\alpha = 0.05$ | 4.3% | 3.9% | +0.4 pp |
| family recovery, known species (LSO) | **91.4%** | 72.2% | +19.2 pp |
| family recovery, novel genus (LGO) | **52.6%** | 28.2% | +24.4 pp |

$n = 1,632$ calibration knowns, 1,632 test knowns and 4,444 novel-genus queries, identical for both scores.
The AUROC difference is +0.065 (95% CI +0.046 to +0.085, 10,000 paired bootstrap resamples clustered on query genus in both classes). The DeLong test on the same contrast gives $z = 11.9$ , $p < 10^{-16}$ ; it assumes within-class independence, which the LGO design violates, so the clustered interval is the one to read. Clustering widens the interval by 1.8 × . McNemar on the $\alpha = 0.05$ flag sets separates power from calibration: among novel queries, 872 were flagged by identity alone against 117 by cosine alone ( $p < 10^{-16}$ ), while the observed false-novelty rates differ by 0.4 percentage points.
619 of the 4,444 LGO queries (13.9%) returned no VSEARCH hit at identity $\geq 0.5$ and query coverage $\geq 0.8$ and carry the censoring value, against 542 under the default-settings search. All are flagged at $\alpha = 0.05$ : of the 1,251 novel queries identity flags there, 619 (49%) are censored rather than scored, so about half of identity's detections on this benchmark are queries with no alignable reference.
The M4 cosine column comes from the leakage-safe retraining, fitted to 11% fewer records and 483 fewer genera than primary M4. DEV was not reopened for that run, so the cost of the mask to model quality is unmeasured.
The calibration and test halves of the LSO set are drawn by `np.random.default_rng(0).permutation`. Across ten halvings (seeds 0 to 9) the identity advantage ranged from +0.063 to +0.075 AUROC and every genus-clustered interval excluded zero; seed 0, reported here, lies near the low end. The operating points should therefore be read as approximate.
The currently deposited splitter yields 3,264 LSO queries rather than the 3,170 of the earlier novelty manuscript; the LGO count reproduces exactly. These results are therefore a paired comparison on the currently reproducible harness, not a numerical reproduction of the earlier LSO run.

**Figure S4:**
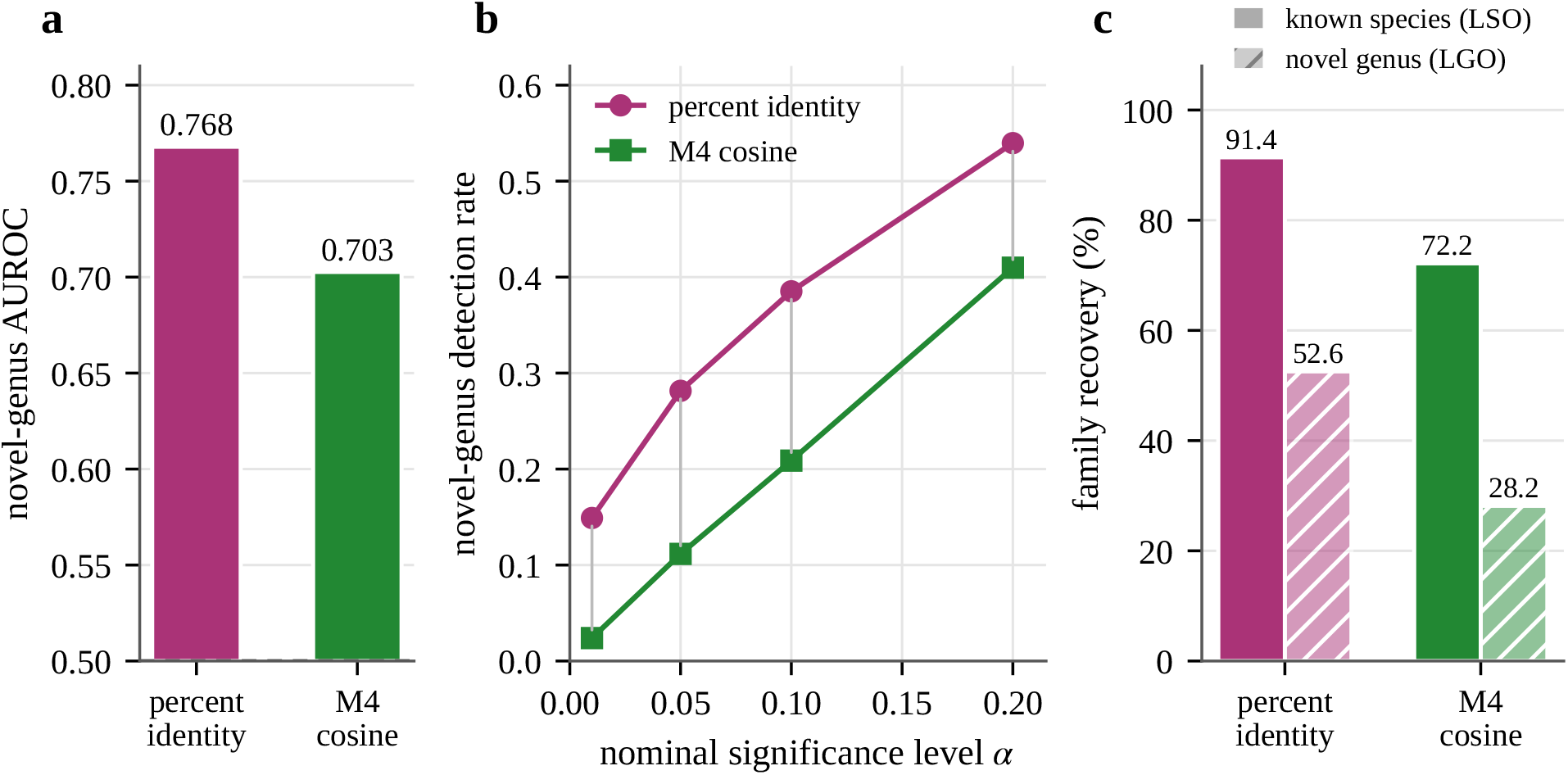
Historical ITS2 benchmark: exhaustive, coverage-filtered identity against leakage-safe M4 cosine with exact-length inference. (a) Novel-genus AUROC. (b) Detection across conformal levels. (c) Family recovery.

## References

[1] Angelopoulos, A.N., Bates, S. (2023). Conformal prediction: a gentle introduction. Foundations and Trends in Machine Learning 16(4), 494–591. doi:10.1561/2200000101

[2] Badirli, S., Akata, Z., Mohler, G., Picard, C., Dundar, M.M. (2021). Fine-grained zero-shot learning with DNA as side information. In Advances in Neural Information Processing Systems 34, 19352–19362.

[3] Badirli, S., Picard, C.J., Mohler, G., Richert, F., Akata, Z., Dundar, M. (2023). Classifying the unknown: insect identification with deep hierarchical Bayesian learning. Methods in Ecology and Evolution 14(6), 1515–1530. doi:10.1111/2041-210X.14104

[4] Bengtsson-Palme, J., Ryberg, M., Hartmann, M., et al. (2013). Improved software detection and extraction of ITS1 and ITS2 from ribosomal ITS sequences of fungi and other eukaryotes for analysis of environmental sequencing data. Methods in Ecology and Evolution 4(10), 914–919. doi:10.1111/2041-210X.12073

[5] Fujisawa, T., Imai, T. (2026). Performance and limitations of out-of-distribution detection for insect DNA barcoding. Ecology and Evolution. doi:10.1002/ece3.73112

[6] Hendrycks, D., Gimpel, K. (2017). A baseline for detecting misclassified and out-of-distribution examples in neural networks. In International Conference on Learning Representations. arXiv:1610.02136.

[7] Ji, Y., Zhou, Z., Liu, H., Davuluri, R.V. (2021). DNABERT: pre-trained bidirectional encoder representations from transformers model for DNA-language in genome. Bioinformatics 37(15), 2112–2120. doi:10.1093/bioinformatics/btab083

[8] Khosla, P., Teterwak, P., Wang, C., Sarna, A., Tian, Y., Isola, P., Maschinot, A., Liu, C., Krishnan, D. (2020). Supervised contrastive learning. In Advances in Neural Information Processing Systems 33, 18661–18673.

[9] Nilsson, R.H., Larsson, K.-H., Taylor, A.F.S., et al. (2019). The UNITE database for molecular identification of fungi: handling dark taxa and parallel taxonomic classifications. Nucleic Acids Research 47(D1), D259–D264. doi:10.1093/nar/gky1022

[10] O’Brien, A., Marín, C., Parada, P. (2026). A calibrated novelty flag for fungal ITS metabarcoding: choosing the error rate at which sequences are declared new. bioRxiv 2026.07.29.741524. doi:10.64898/2026.07.29.741524

[11] O’Brien, A., Marín, C., Parada, P. (2026). Pretrained deep-learning ITS classifiers read the flanking regions, not the ITS2 barcode, and so fail on the amplicon that environmental fungal surveys sequence. bioRxiv 2026.07.29.741510. doi:10.64898/2026.07.29.741510

[12] van den Oord, A., Li, Y., Vinyals, O. (2018). Representation learning with contrastive predictive coding. arXiv 1807.03748.

[13] Romeijn, L., Bernatavicius, A., Vu, D. (2024). MycoAI: fast and accurate taxonomic classification for fungal ITS sequences. Molecular Ecology Resources 24(8), e14006. doi:10.1111/1755-0998.14006

[14] Rognes, T., Flouri, T., Nichols, B., Quince, C., Mahé, F. (2016). VSEARCH: a versatile open source tool for metagenomics. PeerJ 4, e2584. doi:10.7717/peerj.2584

[15] Snell, J., Swersky, K., Zemel, R. (2017). Prototypical networks for few-shot learning. In Advances in Neural Information Processing Systems 30, 4077–4087.

[16] Sun, Y., Ming, Y., Zhu, X., Li, Y. (2022). Out-of-distribution detection with deep nearest neighbors. In Proceedings of the 39th International Conference on Machine Learning, PMLR 162, 20827–20840.

[17] Abarenkov, K., Zirk, A., Piirmann, T., Pöhönen, R., Ivanov, F., Nilsson, R.H., Kõljalg, U. (2025). UNITE general FASTA release for Fungi. Version 19.02.2025. UNITE Community. doi:10.15156/BIO/3301229

[18] Vaze, S., Han, K., Vedaldi, A., Zisserman, A. (2022). Open-set recognition: a good closed-set classifier is all you need? In International Conference on Learning Representations.

[19] Vovk, V., Gammerman, A., Shafer, G. (2005). Algorithmic Learning in a Random World. Springer.

[20] Wang, Q., Garrity, G.M., Tiedje, J.M., Cole, J.R. (2007). Naive Bayesian classifier for rapid assignment of rRNA sequences into the new bacterial taxonomy. Applied and Environmental Microbiology 73(16), 5261–5267. doi:10.1128/AEM.00062-07

[21] Abarenkov, K., Somervuo, P., Nilsson, R.H., Kirk, P.M., Huotari, T., Abrego, N., Ovaskainen, O. (2018). PROTAX-fungi: a web-based tool for probabilistic taxonomic placement of fungal internal transcribed spacer sequences. New Phytologist 220(2), 517–525. doi:10.1111/nph.15301

[22] DeLong, E.R., DeLong, D.M., Clarke-Pearson, D.L. (1988). Comparing the areas under two or more correlated receiver operating characteristic curves: a nonparametric approach. Biometrics 44(3), 837–845.

[23] Edgar, R.C. (2016). SINTAX: a simple non-Bayesian taxonomy classifier for 16S and ITS sequences. bioRxiv. doi:10.1101/074161

[24] Edgar, R.C. (2018). Accuracy of taxonomy prediction for 16S rRNA and fungal ITS sequences. PeerJ 6, e4652. doi:10.7717/peerj.4652

[25] Orsholm, J., Zito, A., Somervuo, P., Harrison, J.P., Koskela, M., Ovaskainen, O., Braga, M.P., Chazot, N., Roslin, T., Furneaux, B. (2026). Discovering the unseen: a performance comparison of taxonomic classification methods for unknown DNA barcodes. Methods in Ecology and Evolution. doi:10.1111/2041-210x.70358

[26] Zito, A., Rigon, T., Dunson, D.B. (2023). Inferring taxonomic placement from DNA barcoding aiding in discovery of new taxa. Methods in Ecology and Evolution 14(2), 529–542. doi:10.1111/2041-210X.14009

